# Impact of aging on the immunological and microbial landscape of the lung during non-tuberculous mycobacterial infection

**DOI:** 10.1101/2023.02.11.528140

**Authors:** Isaac R. Cinco, Nicholas S. Rhoades, Ethan G. Napier, Michael Davies, Derek B. Allison, Steven G. Kohama, Luiz Bermudez, Kevin Winthrop, Cristina Fuss, Eliot R. Spindel, Ilhem Messaoudi

**Affiliations:** Department of Microbiology, Immunology, and Molecular Genetics, College of Medicine, University of Kentucky, Lexington, KY, USA; Division of Neuroscience, Oregon National Primate Research Center, Oregon Health & Science University, Beaverton, OR, USA; Department of Pathology & Laboratory Medicine, College of Medicine, University of Kentucky, Lexington, KY, USA; Department of Microbiology, College of Sciences, Oregon State University, Corvallis, OR, USA; Division of Infectious Diseases, Schools of Medicine and Public Health, Oregon Health and Sciences University, Portland, OR, USA; Department of Diagnostic Radiology, School of Medicine, Oregon Health & Science University, Portland, OR, USA

**Author notes:** Contributed equally to the work (Isaac R. Cinco, Nicholas S. Rhoades).

**Keywords:** Lung, Pulmonary, Aging, non-tuberculous mycobacteria, *M. avium* complex, immunity, microbiome, dysbiosis, rhesus macaque.

## Abstract

Nontuberculous mycobacteria (NTM) are environmentally ubiquitous and predominately cause pulmonary disease (NTMPD). The incidence of NTMPD has steadily increased and is now more prevalent than that of *Mycobacterium tuberculosis* (*M. tb*) in the US. Moreover, the prevalence of NTMPD increases with age; therefore, it is likely that the burden of NTMPD will continue to increase in the coming decades as the number of those over the age of 65 increased in the U.S population. However, the mechanisms leading to higher susceptibility and severity of NTMPD with aging are poorly defined. Here, we used a rhesus macaque model of intrabronchial infection with *M. avium* complex in young and aged animals to address this knowledge gap. Unilateral infection resulted in a robust inflammatory response predominantly in the inoculated lung, however, immune cell infiltration and antigen-specific T cell responses were detected in both lungs. Nasal, oral, and fecal swabs, and BAL samples were profiled using 16S amplicon sequencing. These data suggested that decompartmentalization of the lower respiratory microbiome was occurring, evidenced by detection of bacterial DNA typically found in the gut and oral-pharyngeal cavity in bronchoalveolar samples following infection. Radiographic studies, gross pathology, and histopathology examination revealed increased disease severity in aged compared to young animals with pulmonary consolidation, edema, and lesions. Finally, single cell RNA sequencing indicated that aged animals generated a dysregulated macrophage and CD8 T cell response to MAC infection.

## INTRODUCTION

Nontuberculous mycobacteria (NTM) pulmonary disease (NTMPD) is caused by environmentally ubiquitous non-tuberculous mycobacteria, notably *Mycobacterium avium* complex (MAC; composed of *M. avium* and *M. avium intracellulare*), *M. abscessus*, and *M. kansasii* [1–6]. These bacteria reside in soil and water sources [7], and are inhaled by individuals when aerosolized [8]. Despite the ubiquitous distribution of these bacteria, only a small subset of individuals develop NTMPD, primarily older adults, those with cystic fibrosis (CF), or underlying lung conditions such as COPD [5, 9]. NTMPD is characterized by coughing, shortness of breath, exhaustion, weight/appetite loss, and mild fever. The incidence of this disease has increased from 12 cases per 100,000 persons in 2008 to 18 cases per 100,000 persons in 2015 [10, 11]. Most of these cases are from the elderly population since disease is extremely rare in healthy individuals under 40 years old. Indeed, incidence was reported to be five times higher in patients over 80 years old [4]. In addition, the incidence of NTMPD has recently exceeded that of *Mycobacterium tuberculosis* (*M. tb*) in the United States [5]. Given the increase in the number of older Americans, the incidence of NTMPD is likely to significantly increase over the next decades [12, 13].

Recent studies have suggested a role for the respiratory microbiota in modulating NTMPD susceptibility as evident from the reported differences in the microbiome of the healthy lung versus the NTM infected lung [14–16]. A healthy respiratory microbiome is generally dominated by microbes from the phyla Firmicutes, Bacteriodetes, Proteobacteria, Actinobacteria, and Fusobacteria [14, 17]. Healthy aging was associated with increased abundance of Firmicutes and decreased abundance of Proteobacteria [17]. Comparison of sputum samples from healthy women and those with NTMPD revealed a decrease in *Leptotrichia*, *Streptococcus*, and *Veilonella* [15]. Another study showed that the microbiome of patients with NTMPD sampled by brochoalveolar lavage (BAL) is enriched in *Ralstonia*, *Clavibacter*, *Enterobacter*, and *Mycobacterium* compared to patients with non-CF bronchiectasis [16]. Although these studies provide information about the dysbiosis observed in NTMPD patients, there are no data describing a microbiome that increases risk of NTMPD; therefore, more studies are needed to determine whether alterations in the lung microbial community is a driving factor or simply a symptom of NTM infection.

Risk factors for developing NTMPD include chronic lung diseases such as emphysema or bronchiectasis, the incidence of which increases with age thereby providing a potential explanation for increased rates of NTMPD in older individuals [18, 19]. Additionally, aging is accompanied with several structural and immunological changes in the lung that may contribute to increased risk of acquiring NTMPD such as dilation of air spaces, increase in alveolar dead space, and reduced frequency of macrophages [20]. Moreover, several clinical and genome-wide association studies (GWAS) suggest that T-helper 1 (Th1) T cell responses are necessary for an effective immune response against NTM [21]. Indeed, several mutations in genes important for Th1 response are associated with increased risk of NTMPD in children (*IL12B*, *IL12RB1*, *ISG15*, *IFNGR1*, *IFNGR2*, *STAT1*, and *IRF8*) [22–25] and anti-TNFα therapies increase risk of NTMPD [26, 27]. It has also been shown that peripheral blood mononuclear cells (PBMCs) from patients with NTMPD produced lower levels of IFNγ and TNFα compared to those of healthy controls [27, 28]. Finally, human genetics studies have revealed that toll-like receptor (TLR) signaling, and innate immunity are necessary to combat disseminated NTM [29–34].

Studies carried out using immune competent rodents have provided important insight into the pathogenesis of disseminated NTM infection [35, 36]. A study utilizing an aerosol route of MAC infection in a modified Cornell-like murine model identified IFN as a critical cytokine for the defense against disseminated NTM disease [37]. However, disseminated NTM is largely different from NTMPD where infection is often limited to the middle lobe of the right lung [38, 39], making it unclear whether these findings are relevant in a pulmonary disease setting. Additionally, the specific pathogen free status (SPF) of rodents results in considerable differences in the immunological and microbial respiratory landscapes [40–44] compared to humans who are constantly exposed to pathogens and irritants which modulate lung-resident immune cells and the microbial community throughout the lifespan.

To overcome these hurdles and gain more insight in lung immunological and microbial landscape during NTMPD, we leveraged a rhesus macaque nonhuman primate model of M. avium infection [45]. Rhesus macaques share significant homology with human respiratory and immune systems and recapitulate the hallmarks of human aging [45–47]. In addition, similar to humans, the lung microbiome of healthy rhesus macaques is predominantly composed of microbes from the phyla Actinobacteria (*Tropherema*), Firmicutes (*Streptococcus* and *Veillonella*), Bacteriodetes (*Prevotella*), Proteobacteria (*Neisseria*), and Fusobacteria (*Fusobacterium*) [48]. We previously showed that intrabronchial inoculation of rhesus macaques with MAC resulted in the development of bronchiectasis and formation of granulomatous lesions in the lung [45]. Moreover, the presence of granulomas was associated with a dampened Th1 response indicated by lower frequencies of IFNγ+ T cells and a Th2 bias evidenced by higher levels of antibodies and IL-6 cytokines [45]. In the current study, young and aged rhesus macaques were inoculated intrabronchially with MAC into the right caudal lung. Radiological changes, host responses and composition of microbial community in the lung were evaluated longitudinally post challenge. Overall, our findings suggest that NTM infection results in more severe disease in aged animals that was accompanied by alterations in both innate and adaptive immune responses as well as significant shifts in the lung microbial community indicative of decompartmentalization.

## MATERIALS AND METHODS

### Ethics Statement

Animal work was performed in strict accordance with the recommendations described in the Guide for the Care and Use of Laboratory Animals of the National Institute of Health, the Office of Animal Welfare and the Animal Welfare Act, United States Department of Agriculture. The studies were approved by the Institutional Animal Care and Use Committees (IACUC) at the Oregon National Primate Research Center. Procedures were conducted in animals anesthetized by trained personnel under the supervision of veterinary staff. All efforts were made to enhance animal welfare and minimize animal suffering in accordance with the Weatherall report [49] on the use of nonhuman primates in research.

### Cohort description, animal infection, and sample collection

Three young (5-10 years) and two aged (>18 years) colony-bred Indian-origin rhesus macaques (*Macaca mulatta*) were intrabronchially inoculated in the right caudal lung with 6.8×10^8^ CFU of *M. avium* subsp. *hominissuis* strain 101 (MAC) (**Supp. Table 1**). Blood, bronchoalveolar lavage (BAL), and oral, nasal, and rectal swabs were collected on days 0, 8, 15, 28, 44, 56, 86, 121, 149, and 190 post infection. Animals underwent a chest computed tomography (CT) scan every 3-4 weeks. Animals were euthanized at 190 days post infection (DPI).

### Bacterial culture and load

M. avium complex subsp. *hominissuis* strain 101, available at ATCC, was obtained from from a patient with AIDS and shown to be virulent in mouse models, including a mouse model of pulmonary infection [35]. The bacteria were obtained from a from stock and grown on Middlebrook 7H10 agar. An inoculum was established (approximately 1 x 10^6^ bacteria prior to infection). The viability of the inoculum was 94 +/− 2% determined as previous described [35]. Bacterial load was determined by enumerating the number of colony-forming units (CFU) in BAL supernatant after 8 weeks of incubation at 37°C/5% CO_2_ onto Lowenstein-Jensen agar plates and real-time quantitative PCR (qPCR). DNA was extracted from BAL cells using the DNeasy PowerSoil Pro Kit (Qiagen, Germantown, MD) according to the manufacturer’s instructions. Bacterial burden of MAC at the time points previously listed was determined by quantitative real-time PCR (qPCR) using TaqMan Universal PCR Master Mix (Thermo Fisher Scientific, Waltham, MA) and primers/probe specific for the IS1311 insertion sequence found in *M. avium* [50]. Each run was initiated at 50°C for 2 min, then 95°C for 10 min, followed by 40 cycles at 95°C for 15 sec then 60°C for 1 min using a QuantStudio 3 Real-Time PCR System (Thermo Fisher Scientific). Extracted MAC DNA was used as the quantification standard.

### Electron Microscopy

Swabs of the airways and intra-airway plug were washed twice, and the pellet was suspended in fixative buffer with 2.5% glutaraldehyde, 1% formaldehyde and 0.1 M sodium cacodylate. Sections were stained, dehydrated, and visualized for SEM and TEM in the EM facility of Oregon State University as reported [51].

### Flow Cytometry

1 x 10^6^ BAL cells were surface stained with antibodies against CD20, CD27, IgD, CD4 (Biolegend, San Diego, CA), CCR7 (BD Biosciences, Franklin Lakes, NJ), CD8b (Beckman Coulter, Brea, CA), and CD28 (Tonbo Biosciences, San Diego, CA). Cells were then fixed and permeabilized before the addition of anti-Ki67 (BD biosciences, Franklin Lakes, NJ) to assess proliferation. This panel allows the identification of B cells (CD20) and T cells (CD8b, CD4) as well as naïve and memory subsets as previously described [48]. Another 10^6^ BAL cells were stained with CD206, CD14, HLA-DR, CD8a, CD123, CD16 (Biolegend, San Diego, CA), CD11c (Invitrogen, Waltham, MA), and Granzyme-B (BD Biosciences) to assess frequency of innate immune cells [48]. To measure response of antigen-specific T cells, monocyte-derived macrophages, and DCs after MAC infection, 10^6^ BAL cells were stimulated with 1 mg/mL MAC lysate in the presence of brefeldin A (BFA) for 16 hrs. At the end of the incubation, cells were surface stained with CD8b (Beckman Coulter), CD4, CD20, CD14, HLA-DR (Biolegend); cells were then fixed and permeabilized followed by staining with IL-17, IFNγ (Biolegend), and TNFα (Invitrogen). Samples were acquired using Attune NxT (Life Technologies, Carlsbad, CA) and analyzed with FlowJo software (TreeStar, Ashland, OR).

### Luminex

Levels of immune mediators in BAL supernatant were analyzed using the R&D 36-plex NHP XL Cytokine Premixed Kit (Bio-Techne, Minneapolis, MN). The following analytes were measured: BDNF, CCL2, CCL5, CCL11, CCL20, CD40L, CXCL2, CXCL10, CXCL11, CXCL13, FGF basic, G-CSF, GM-CSF, Granzyme B, IFNα, IFNβ, IFNγ, IL-1β, IL-10, IL-12 p70, IL-13, IL-15, IL-17A, IL-2, IL-21, IL-4, IL-5, IL-6, IL-7, IL-8, PDGF-AA, PDGF-BB, PD-L1, TGFα, TNFα, VEGF. Samples were acquired using the MAGPIX xMAP (Luminex Corporation, Austin, TX). Data were analyzed using the Luminex Xponent software with an 8 point logistic regression curve.

### ELISA

IgG antibody titers against MAC were determined using enzyme-linked immunosorbent assay (ELISA). Plates were coated with 1 mg/mL MAC bacterial lysate overnight at 4°C. The plates were washed three times with 0.05% Tween-PBS. Heat-inactivated (56°C for 30 min) plasma samples were then added in 3-fold dilutions in duplicates for 1.5 hr at room temperature. After washing with 0.05% Tween-PBS, Goat anti-monkey IgG (Fc) HRP (Brookwood Biomedical, Jemison, AL) was diluted then added to each well and incubated for 1-1.5 hr. The plates were then washed three times with 0.05% Tween-PBS and incubated for 20 min in a solution containing o-phenylenediamine·2HCl (OPD) substrate (Sigma-Aldrich, St. Louis, MO) diluted in substrate buffer and 30% H_2_O_2_. The reaction was stopped by adding 1M HCl to each well. Absorbance at 490 nm was measured using the SpectraMax iD3 (Molecular Devices, San Jose, CA) plate reader. Each plate included a positive control which was used to normalize the data.

### Immunohistochemistry and Immunofluorescence analysis

Anti-CD3 (T cells), anti-CD20 (B cells), anti-CD68/CD163 (monocyte/macrophage), and anti-myeloperoxidase antibodies were used to stain lung tissue from young and aged rhesus macaques. Whole slide multispectral digital images were processed utilizing the HALO Image Analysis platform (version X, Indica Labs). Image analysis algorithms were constructed using the High-Plex FL module (v4.1.3). DAPI was utilized for nuclear segmentation, and unique thresholds for cytoplasmic positivity were set for each antibody.

### 16s Amplicon Sequencing and Bioinformatics Analysis

Amplification of the hypervariable V4 region of the 16s rRNA gene was performed using the 515F/806R PCR primers; the forward primers were conjugated with a 12-bp barcode [52]. Each reaction was run in duplicate and prepared with GoTaq master mix (Promega Corporation, Madison, WI) according to manufacturer’s instructions. Cycling conditions were: 94°C for 3 min, 37 cycles of 94°C for 45 s, 50°C for 1 min, and 72°C for 1 min, followed by a final extension at 72°C for 10 min. The PCR products were multiplexed using Quant-iT PicoGreen dsDNA Assay Kits and dsDNA Reagents (Thermo Fisher). The resulting library was then spiked with ∼15-20% PhiX and sequenced on an Illumina MiSeq. Raw FASTQ 16s rRNA gene amplicon sequences were processed using the QIIME2 analysis pipeline [53]. Sequences were demultiplexed and filtered using DADA2, resulting in the removal of chimeric sequences and the generation of an amplicon sequence variant (ASV) table [54]. MAFFT was used to align the sequence variants while FastTree2 was utilized to construct a phylogenetic tree [55, 56]. Taxonomy was assigned to sequence variants using q2-feature-classifier against the SILVA database (release 138) [57]. The samples were rarified to 10,000 sequences per sample to normalize the sequencing depth between samples. QIIME2 was also used to generate alpha diversity metrics, which include richness (as observed ASV), Shannon evenness, and phylogenetic diversity, while beta diversity was estimated using weighted and unweighted UniFrac distances [58]. The LEfSe algorithm allowed for the identification of differentially abundant bacteria between groups with a linear discriminant analysis score cutoff of 2 [59].

### Single Cell RNA library generation and bioinformatics analysis

Cryopreserved BAL cells from 0, 8, 14, and 86 DPI were thawed then stained with Sytox orange (Thermo Fisher) to sort viable cells using an iCyt-Sony Cell Sorter System. Live cells were counted on a TC20 Automated Cell Counter (BioRad), tagged with cell multiplexing oligos (CMOs) to allow for downstream multiplexing, and pooled. The cells within each pool were diluted to a concentration of 1600 cells/μL in ice-cold PBS with 0.04% BSA. Single cell suspensions were loaded on the 10X Genomics Chromium Controller with a loading target of 20,000 cells. The libraries were prepared using the Chromium Single Cell 3’ Feature Barcoding Library Kit with the v3.1 chemistry (10X Genomics, Pleasanton, CA). Libraries were sequenced using an Illumina NovaSeq with a target of 30,000 reads per cell RNA library and 2000 reads per cell hashtag-oligo barcode library.

The reads were aligned and quantified using the Cell Ranger Single-Cell Software Suite (version 4.0, 10X Genomics) against the Mmul_8 rhesus macaque reference genome using the STAR aligner. Seurat (version 4.1.1) was used for downstream analysis of the aligned reads. The libraries were merged and droplets with ambient RNA (<200 feature counts) and dying cells (>20% total mitochondrial gene expression) were filtered [60]. The resulting multiplexed library was further processed by performing variance stabilization with the *SCTransform* function, which generates a regularized negative binomial regression corrected for differential effects of mitochondrial and ribosomal gene expression levels, followed by data normalization using *NormalizeData*. The data was then scaled using *ScaleData* in preparation for PCA generation which was accomplished using *RunPCA*. This allowed us to determine dimensionality of our sample i.e., 30 principal components (PC) were used to cluster the cells before UMAP generation using the *FindNeighbors* and *FindClusters* (resolution = 0.5) function in Seurat. Finally, Seurat’s *runUMAP* function allowed for UMAP generation. Cell types were assigned to individual clusters using *FindMarkers* function with a Log2 fold change cutoff of 0.4. Differential expression analysis was performed using MAST with default settings in Seurat. All comparisons were performed between day 0 and days of notable DPI (8,14, and 86). Gene expression changes were considered as significant with a log fold change ≥ 0.58 and p ≤ 0.05.

### Statistical Analysis

Statistical analysis was performed using the GraphPad Prism software (GraphPad Software Inc., La Jolla, CA). Statistical significance when comparing 3 groups or more was determined using a 1-way analysis of variance (ANOVA) with the Benjamini and Hochberg false discovery rate (FDR) multiple comparisons method to accommodate the small sample size (n=5) of the study. Significance for the comparison between 2 groups was measured using a two-tailed, unpaired parametric Welch’s T-test.

## RESULTS

### Aged Rhesus macaques exhibit more severe NTMPD following MAC infection

Three young adult (2 females and 1 male) and two aged female rhesus macaques were infected intrabronchially in the right lower lobe of their lung with 6.8×10^8^ CFU of *M. avium* subsp. *hominissuis* strain 101 (MAC) at day 0 (**Figure 1A**). Broncho-alveolar lavage (BAL) samples were collected separately from the right and left lungs to characterize the kinetics and magnitude of the immune responses as well as longitudinal changes in the MAC bacterial burden and lung microbiome. Live bacteria were only detected in the right BAL supernatant 8 days post-infection (DPI) (**Figure 1B**). Similarly, bacterial burden measured by real-time quantitative PCR (qPCR) peaked within the right lungs at 8 DPI followed by a rapid decline (**Figure 1B**). Bacterial DNA in the left lung was only sporadically detected in aged 1 (A1) and young 1 (Y1), indicating limited movement of the bacteria from the site of the inoculation (**Figure 1B**). Conversely, scanning electron microscopy revealed that *M. avium* was a part of biofilms in both the left and right lungs (**Supp. Figure 1A, B**). An aggregate of host cells (plug) found in the right lung exhibited tick secretions (**Supp. Figure 1C**). Transmission electron microscopy of an alveolar macrophage in the airway plug showed that there were viable *M. avium* present within the cell (**Supp. Figure 1D**).

**Figure 1:**
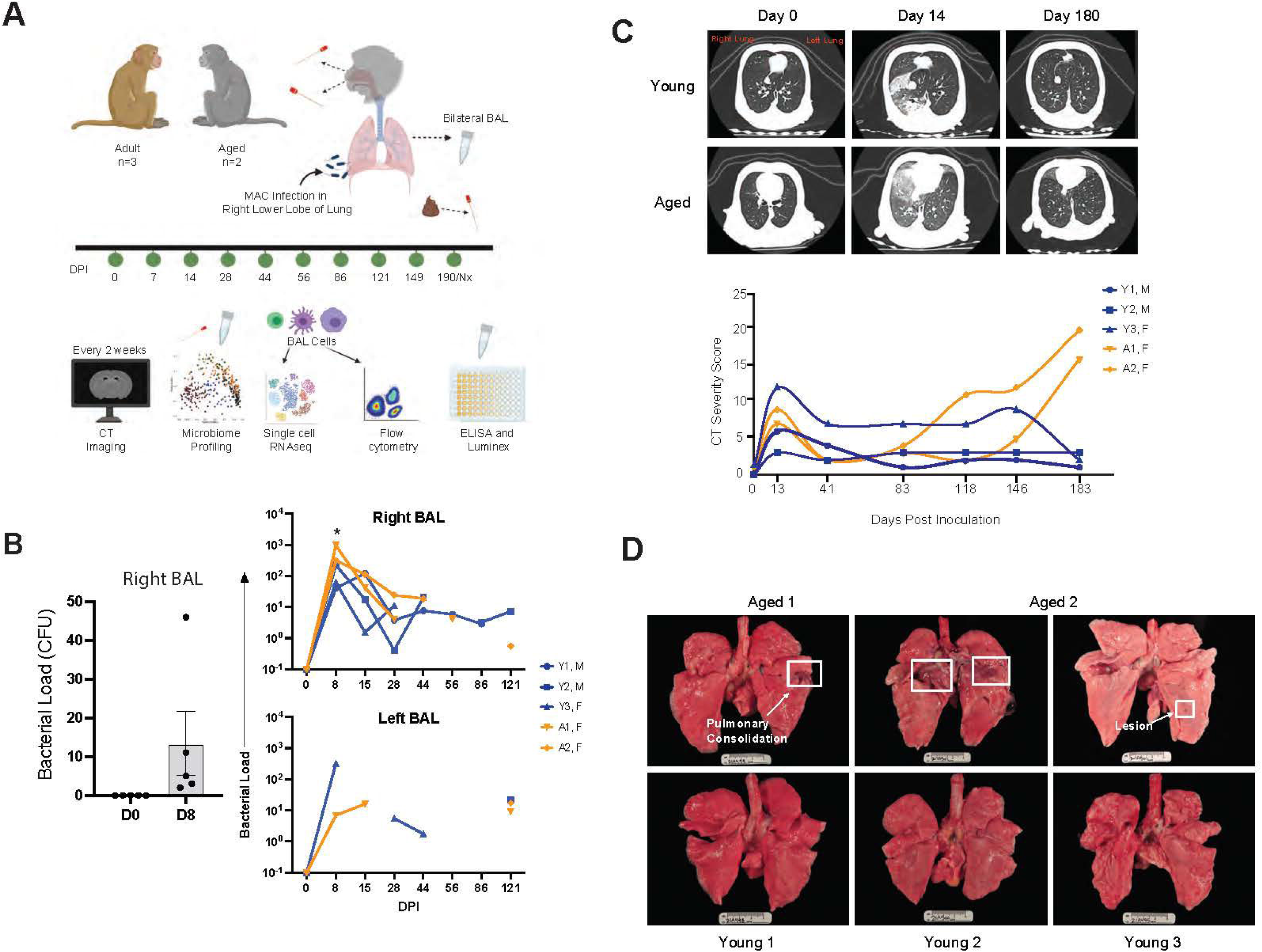
Age-related differences of NTMPD progression are observed in rhesus macaques. **(A)** Experimental design. Three young adult (5-10 years) and two aged (>18 years) rhesus macaques were inoculated in the right caudal lung with a bolus of 6.8×10^8^ CFU of MAC. Nasal, oral, fecal, and BAL samples were collected at the indicated days post inoculation (DPI). BAL cells were probed via microbiome profiling, scRNA-seq, flow cytometry, ELISA, and Luminex assays. Microbiome profiling was done on the nasal, oral, and fecal swabs. Computed tomography (CT) scans and severity scores were collected every two weeks. **(B)** Bacterial load was measured in the right and left BAL supernatants via colony forming units (CFU) after 8 weeks of incubation onto Lowenstein-Jensen agar plates and quantitative real-time PCR specific for the IS1311 sequence found in *M. avium*. Missing data points were those that had a quantification cycle (Cq) value below the threshold. Data for all animals were compared at each timepoint to 0 DPI (*p < 0.05). **(C)** Representative CT scans of the young and aged animals before inoculation (0 DPI), at the height of inflammation (14 DPI), and close to necropsy (180 DPI). CT severity scores were plotted at the indicated days post inoculation. **(D)** Gross lung pathology exhibiting pulmonary consolidations and lesions in the aged animals compared to the young animals at necropsy. Data from young animals (Y1-Y3) plotted in blue, aged animals (A1, A2) plotted in orange. M indicates male animals and F indicates females.

Each animal underwent computed tomography (CT) scans to assess longitudinal radiographic changes in response to infection. In line with the peak in bacterial load at 8 DPI, the CT scans showed considerable inflammation 14 DPI in both young and aged animals (**Figure 1C**). However, while radiographic changes resolved in young animals by 41 DPI, aged animals experienced an exacerbation of inflammatory changes starting at 118 DPI that were unresolved, even out to 190 DPI (necropsy) (**Figure 1C**). In accordance with these observations, signs of pulmonary consolidation and lesions were only evident in the lungs from aged animals (**Figure 1D**). In contrast, no overt pathology was observed in any of the lungs from young animals (**Figure 1D**).

### Increased immune infiltrates detected with age

Histopathology revealed significant cellular and structural changes within the lungs at necropsy (**Figure 2**). Hematoxylin and eosin (H&E) staining of lung sections shows immune infiltrates, lymphoid aggregates, and signs of interstitial fibrosis such as collagen deposition, pulmonary edema, and bronchopneumonia in lungs from aged animals (**Figure 2A-F, Supp. Figure 2**). Notably, in animal A2, pulmonary consolidation with intraalveolar chronic inflammation, interstitial edema, and interstitial fibrosis were observed in the right accessory lobe (**Figure 2A**) while the right caudal lobe showed signs of chronic interstitial and intraalveolar lymphocytes, eosinophils, and neutrophils (**Figure 2B**). Sections of the right caudal lung also showed dense collagen deposition (fibrosis), resulting in interstitial thickening, as well as neutrophil accumulation close to the pleura (**Figure 2B**). The left caudal lung of A2 had a bulla containing neutrophil aggregates (**Figure 2C**). Additionally, areas of consolidation with intersitial edema and intraalveolar inflammation can be seen in the right accessory lobe of A2 (**Figure 2D**). Despite the lack of significant interstitial fibrosis, A1 exhibited chronic interstitial and peribronchiolar inflammation indicated by multiple lymphoid aggregates (**Figure 2E-F**). Finally, Y2 exhibited few lymphoid aggregates and pulmonary edema with a protein rich fluid in the right caudal lobe (**Figure 2G**), however, other sections of this animal’s lungs didn’t show notable histological changes (**Figure 2H, Supp. Figure 2**). The lungs of Y1 and Y3 showed unobstructed airways and pleura, no interstitial fibrosis, and only residual inflammation (**Supp. Figure 2**).

**Figure 2:**
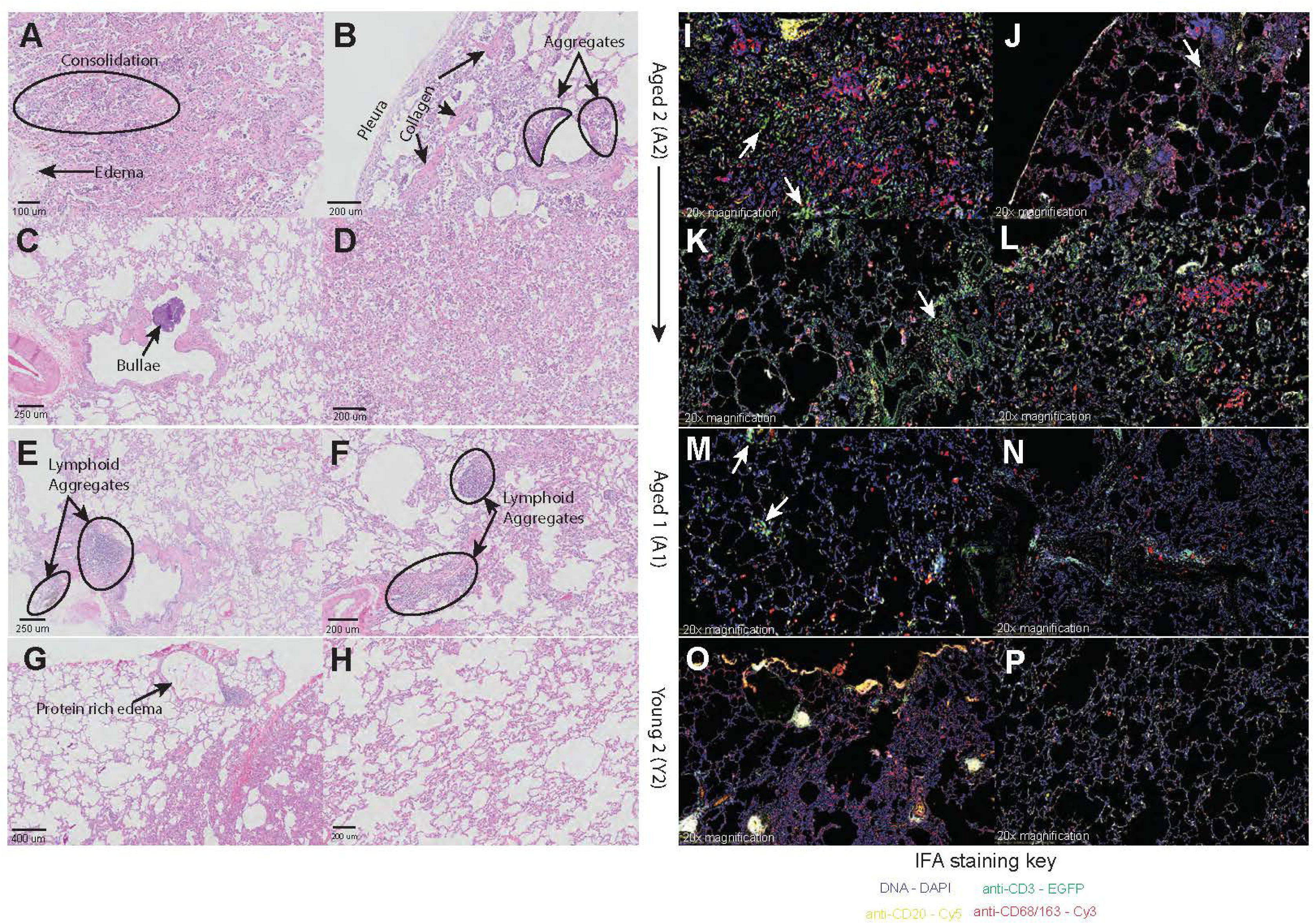
Age-related pulmonary changes and inflammation observed in MAC infected NHP lung samples. H&E stains at the indicated magnifications of aged 2 **(A)** right accessory lobe 1, **(B)** right caudal lobe, **(C)** left caudal lobe, and **(D)** right accessory lobe 2; aged 1 **(E)** right cranial lobe and **(F)** right caudal lobe; young 2 **(G)** right caudal lobe and **(H)** right middle lobe. IFA stains at 20x magnification of aged 2 **(I)** right accessory lobe 1, **(J)** right caudal lobe, **(K)** left caudal lobe, and **(L)** right accessory lobe 2; aged 1 **(M)** right cranial lobe and **(N)** right caudal lobe; young 2 **(O)** right caudal lobe and **(P)** right middle lobe. The fluorescent channels used are 4’,6-diamidino-2-phenylindole (DAPI) stain for a general DNA stain (Blue); enhanced green fluorescent protein (EGFP) conjugated to anti-CD3 for T cell identification (Green); Cyanine 5 (Cy5) conjugated to anti-CD20 for B cell identification (Yellow); Cyanine 3 (Cy3) conjugated to anti-CD68/163 for identification of macrophages and monocytes (Red). White arrows indicate clusters of T or B cells.

Given the histological findings, an immunofluorescence assay (IFA) was performed to better characterize the cellular immune infiltrates (**Figure 2, Supp. Figure 3**). This analysis showed that large numbers of infiltrating T cells were found in the right accessory lobes, right caudal, and left caudal sections in A2 (**Figure 2I-L, Supp. Figure 3**). Similarly, T cell infiltrates as well as T and B cell lymphoid aggregates were detected in A1 (**Figure 2M, N, Supp. Figure 3**). Some lymphoid aggregates can be seen in Y2 (**Figure 2O, Supp. Figure 3**), while all other sections from this and the remaining 2 young animals showed healthy lung tissue (**Figure 2P, Supp. Figure 3**). The immune infiltrates were enumerated (**Supp. Figure 4A-G**) via an algorithm created on the HALO digital image analysis platform utilizing a High-Plex FL module, as described above. This analysis revealed a statistically significant increase in macrophages and T cells in the aged compared to the young animals (**Supp. Figure 4H**).

### The robust inflammatory response is contained within the infected lung

Given the inflammatory changes detected by imaging and histology, we performed a Luminex assay on BAL fluid collected 0-28 DPI to assess changes in immune mediator levels after infection. In the right lung, we observed an increase in the concentration of numerous cytokines (IL-10, IL-12 (p70), IL-17, IFNγ, IFNα, TNFα, CD40L, IL-13, IL-8, IL-6, IL-2) and chemokines (CXCL11, CXCL10, CCL5, CCL20, CXCL13, CXCL2, CCL2) 8-28 DPI (**Figure 3A**). Levels of most immune mediators peaked 8 DPI after which most returned to baseline except for IL-10 and CXCL-10 where the levels remained high 14 and 28 DPI respectively (**Figure 3A**). Levels of growth factors involved in tissue repair and wound healing (PDGF-BB, BDNF, GM-CSF) also increased 8 DPI with GM-CSF remaining significantly elevated 14 DPI (**Figure 3A**). Unlike the other immune mediators, CCL2 (chemokine recruiter of monocytes, basophils, and eosinophils) and VEGF (abundant in lung epithelium and angiogenesis promoter) significantly increased 14 and 28 DPI (**Figure 3A**). We then compared levels of cytokines or chemokines in response to MAC infection between young and aged animals using an area under the curve (AUC) analysis. Modest increases in levels of IL-10 and CD40L were observed in the young compared to aged animals (**Figure 3B**).

**Figure 3:**
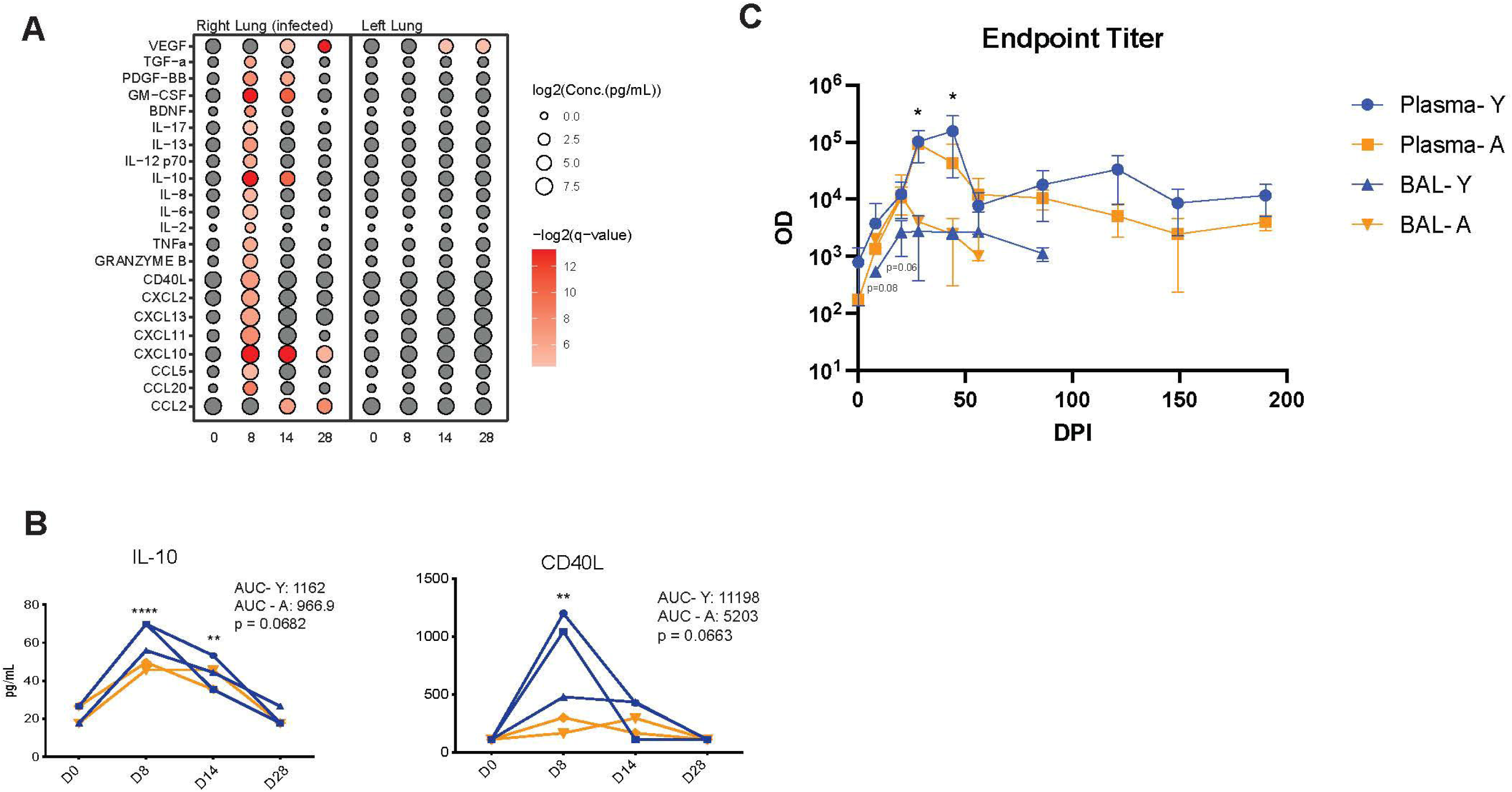
A robust inflammatory response and a modest antibody response occur in the right lung (infected). **(A)** Bubble plot of the most upregulated cytokines as determined by Luminex assays at the indicated timepoints in the right and left lungs. The size of the bubble represents the concentration of the immune mediator while color represents the significance of the change. **(B)** Graphical visualization of the concentration (pg/mL) of IL-10 and CD40L in the right lung. Area under the curve (AUC) was determined for both young and aged groups and compared to each other. **(C)** MAC-specific IgG end point titer averages in the plasma and BAL of the young and aged animals. Each group for all timepoints were compared to 0 DPI. *p < 0.05, **p < 0.01, ***p < 0.001.

Additionally, we performed an ELISA assay on plasma and bilateral BAL supernatants to measure the systemic and local MAC-specific IgG antibody titers, respectively. Plasma IgG levels show a rapid rise from 0-44 DPI followed by a decline and plateau from 56-190 DPI (**Figure 3C**). The BAL IgG profile was similar to the plasma, although, it exhibited a modest and transient increase from 8-14 DPI followed by a steady decline from 14-86 DPI, ultimately reaching nadir levels (**Figure 3C**).

### Robust cellular responses to MAC infection in both the infected right lung and non-infected left lung

To better uncover cellular changes induced by MAC challenge, the frequency and phenotype of immune cells present in a BAL were determined longitudinally during infection by flow cytometry (**Supp Figure 5**). Frequency of the different cell subsets was differentially affected by MAC infection with most significant changes observed in the right lung where the animals were inoculated (**Figure 4**). As expected, prior to infection, alveolar macrophages (AM) were the most abundant immune cells in a BAL. However, at 8 and 14 DPI, the relative frequency of AM declined dramatically in the right lung before rebounding back to pre-infection levels at 44 DPI and fluctuating for the remainder of the study (**Figure 4A**). This early decrease in AM frequency in the right lung was coupled with a significant influx of CD4+ T cells at 8 DPI (**Figure 4B**) while frequencies of CD8 T cell remained unchanged (**Figure 4C**). Frequency of CD4 and CD8 T cells were consistently higher compared to 0 DPI in the left lung (**Figure 4B, C**). Interestingly, the frequency of natural killer (NK) cells increased dramatically in both right and left lung at the later stages of the infection (**Figure 4D**).

**Figure 4:**
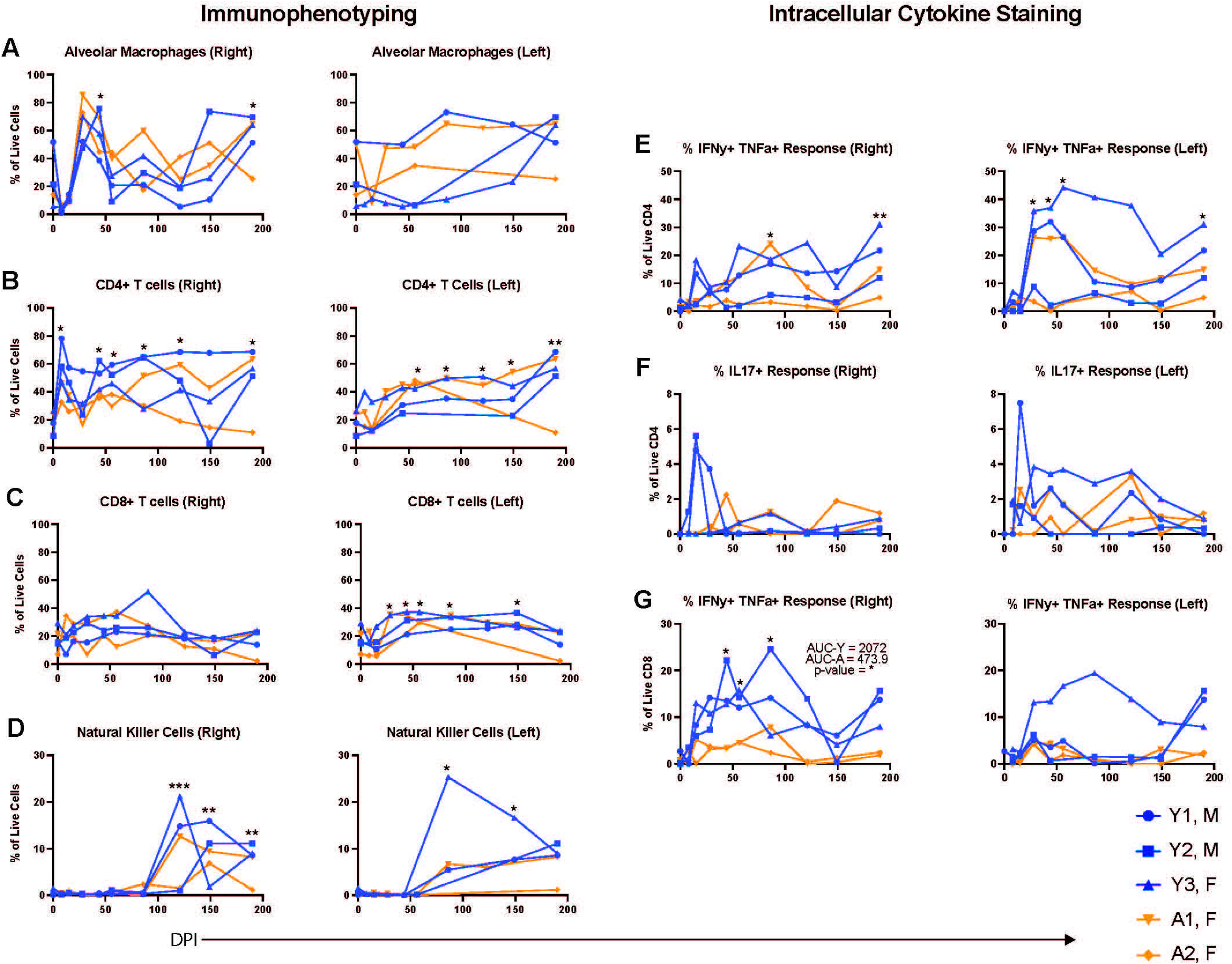
Infiltrating immune cells are observed in both the left and right lung of MAC infected rhesus macaques. Bronchoalveolar lavage (BAL) samples were collected from the right and left lungs at various days post infection (DPI). The percent abundance of live **(A)** alveolar macrophages (AMs), **(B)** CD4+ T cells, **(C)** CD8+ T cells, and **(D)** natural killer (NK) cells are shown. The MAC-specific CD4 and CD8 T cell response was determined by intracellular cytokine staining for IFNγ, TNFα and IL-17 following stimulation with 1 mg/mL MAC lysate. The percent response of **I** IFNγ+ TNFα+ CD4 T cells, **(F)** IL-17+ CD4 T cells, and **(G)** IFNγ+ TNFα+ CD8 T cells are shown. Data from all animals at each timepoint were compared to 0 DPI. Area under the curve (AUC) was determined for both young and aged groups and compared to each other. *p < 0.05, **p < 0.01, ***p < 0.001.

To determine if the infiltrating T cells were antigen-specific, we performed an intracellular cytokine staining (ICS) assay after overnight stimulation with MAC lysate. Although MAC-specific CD4 T cell responses were detected at various time points post infection, large animal-to-animal variation was observed such that a significant increase over baseline was only evident at 86 and 190 DPI in the right lung (**Figure 4E**). In the left lung, however, significantly increased frequency of Mac-specific CD4 T cells was detected 28-56 DPI and at 190 DPI (**Figure 4E**). Only two young animals generated detectable IL-17 CD4 T cell responses in both the right and left BALs (**Figure 4F**). MAC-specific CD8 T cells were detected 44-86 DPI and their frequencies were higher in the young compared to the aged animals in the right BAL (**Figure 4G**). Appreciable CD8 T cell responses were only detected in one young animal in the left BAL (**Figure 4G**).

### Transcriptional changes in the immune landscape of the lungs over the course of infection

To further understand the host response to MAC infection and the impact of age on the immunological response to MAC infection, we profiled BAL cells at four timepoints over the course of infection (DPI 0, 8, 14, and 86) using single-cell RNA sequencing (scRNA-Seq). Data from all the libraries were pooled and subjected to quality check resulting in 79,154 total cells available for downstream analysis. Principal component analysis (PCA) and linear dimensional reduction through uniform manifold approximation and projection (UMAP) revealed several distinct lymphoid and myeloid clusters (**Figure 5A, Supp. Figure 6A**) that were annotated using canonical markers identified using Seurat’s *FindAllMarkers* function (**Figure 5B, Supp. Figure 6B**). AM subsets were identified based on the expression of *FABP4* and *MARCO*, while infiltrating macrophages expressed higher levels of *CD14* and various inflammatory markers, including alarmins (*S100A8*, *S100A9*), complement proteins (*C1QB)*, and pro-inflammatory cytokines *(IL1B*). Dendritic cells (DCs) expressed high levels of *MAMU-DRA* (MHC-II) and *IRF8*. All T cell clusters expressed varying levels of *CD3E* and were delineated into either CD4 or CD8 T cells based on expression of *S100A4*, *CCR6*, and cytotoxicity molecules *GRZMK*, *GRZMB*, and *GNLY* respectively. The NK cells were defined based on the expression of cytotoxic molecules *GNLY*, *GRZM*, and NK cell receptors *KLRD1* in the absence of CD3. B cells exclusively expressed canonical marker *MS4A1* in addition to *CD79B* and *CD79A*. Finally, small clusters of mast cells (*MS4A2*), epithelial cells (*WFDC2* and *CST6*), proliferating T cells (CD3, Ki67) and proliferating AM (FABP4, MARCO, and Ki67) were identified (**Figure 5B, Supp. Figure 6B**).

**Figure 5:**
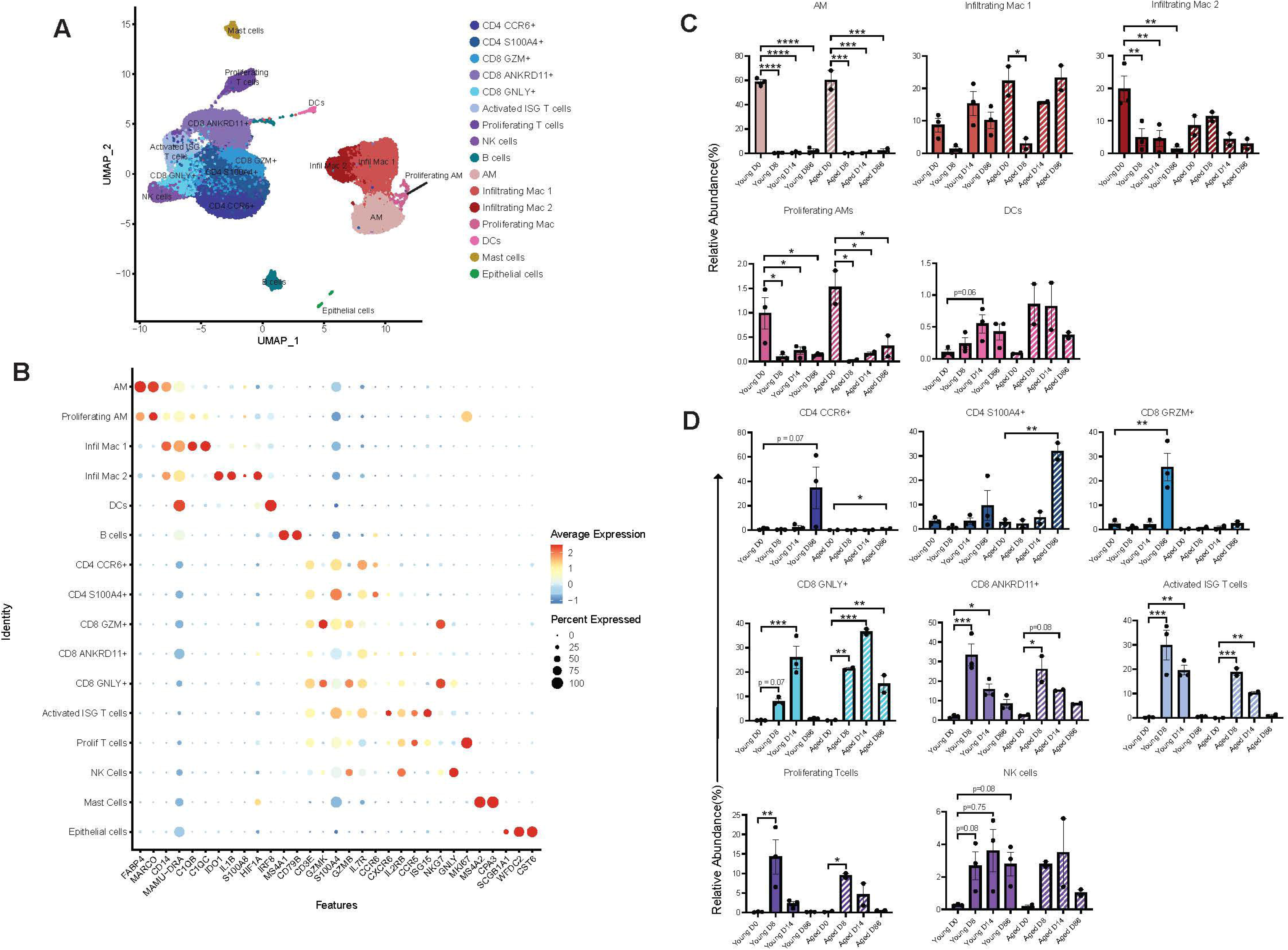
Single cell RNA sequencing analysis reveals shifts in the immune landscape of the lung. **(A)** Uniform manifold approximation and projection (UMAP) representation of 79,154 immune cells within 5 timepoints across MAC inoculation (days 0, 8, 14, 86, 190) showing 16 unique clusters. **(B)** Bubble plot of key gene markers used to annotate the UMAP. **(C)** Significant relative abundances within clusters of myeloid cells by DPI. **(D)** Significant relative abundances within clusters of lymphoid cells by DPI. Significance for panel C and D was calculated using a 1-way ANOVA with Benjamini and Hochberg’s post-hoc FDR comparisons relative to 0 DPI. *p <0.05,**p<0.01,***p<0.001,****p<0.0001.

To further ensure that our classification of myeloid and lymphoid subsets was accurate, we carried out functional enrichment of the marker genes. Myeloid cluster genes enriched to gene ontology (GO) terms associated with inflammation, chemotaxis, and tissue repair (**Supp Figure 6C**). More specifically, marker genes of AM subsets played a role in response to oxidative damage as well as anti-microbial defense, while proliferating AM additionally highly expressed genes associated with cell cycle (**Supp Figure 6C**). Infiltrating macrophages highly expressed genes important for wound healing, chemotaxis, phagocytosis, and inflammation (**Supp Figure 6C**). Finally, marker genes that define DC play a role in endocytosis and regulation of defense response in line with their role in activating T cells (**Supp Fig 6C**).

Markers of CD4 clusters enriched to GO terms in line with functions often attributed to helper T cells such as regulation of immune effector process, immune response, cytokine production, and cell migration (**Supp. Figure 6D**). On the other hand, marker genes of CD8 subsets enriched to GO terms “cytolysis”, “alpha-beta T cell activation”, and “T cell receptor signaling” (**Supp. Figure 6D**). Genes that defined the activated interferon stimulated gene (ISG) T cells enriched to GO terms “response to virus” and “response to type 1 IFN”. The NK cell clusters gene markers play critical roles in cytolysis and C-lectin receptor signaling (**Supp. Figure 6D**). Mast cells expressed genes associated with inflammatory response and wound healing while epithelial cells were enriched to cilium movement and organization (**Supp. Figure 6E**).

We next determined changes in the relative abundance of these clusters throughout infection (**Figure 5C**). As described above using flow cytometry, frequency of AM and infiltrating macrophages in young and old animals decreased sharply 8 DPI. While the frequency of AM and infiltrating macrophage 2 subsets remained low for the rest of the study, the infiltrating macrophage 1 subset rebounded. Frequency of DCs was similar in aged and young with low numbers at 0 DPI followed by a modest increase 14 DPI in the young animals followed by a return to baseline values (**Figure 5C**). Frequencies of all T cell subsets, which were negligible pre-infection, increased following infection in an age and DPI dependent manner (**Figure 5D**). Frequency of CD4 CCR6+ T cells increased modestly in young, but not aged, animals 86 DPI while those of CD4 S100A4+ cells significantly increased at 86 DPI in the aged, but not young animals (**Figure 5D**). The relative abundance of CD8 GLNY+, CD8 ANKRD11+, and ISG T cells increased dramatically 8 and 14 DPI with CD8 GNLY peaking 14 DPI in young and 8 DPI in aged animals but declining thereafter. Proliferating T cells also showed a dramatic increased 8 DPI before returning to pre-infection levels. Finally, frequencies of NK cells increased in young animals 8 DPI and remained increased until 86 DPI (**Figure 5D**).

### Transcriptional shifts in myeloid and lymphoid cells are associated with response to intracellular pathogen

Next, we investigated changes in the transcriptional profiles of the various immune subsets over time. We first measured module scores of gene sets important for “cytokine-chemokine signaling”, “toll-like receptor (TLR) signaling”, and “anti-viral/bacterial responses” (**Supp. Table 2**) longitudinally within all myeloid clusters. Scores for all three modules were significantly increased 8 through 86 DPI in both infiltrating macrophages subsets and proliferating AM (**Figure 6A**). Functional enrichment of differentially expressed genes (DEGs) detected at these time points in the AM subset enriched to “regulation of cellular response to stress” with about 150 genes, or “regulation of neutrophil activation” and “negative regulation of immune system process” with less than 50 genes (**Figure 6B**). Infiltrating macrophage subsets showed over representation of GO terms “regulation of defense response”, “inflammatory response”, and “cellular responses to cytokine stimulus” (**Figure 6C**). In line with these upregulated processes, the expression of several cytokines and chemokines as well as inflammatory genes increased in a time-dependent fashion throughout infection. Expression of several NFκB regulated genes increased significantly at 8 and 14 DPI including *IL6*, *IL1B, NFKBIA, CXCL9, CD86,* and *S100A8*. At 86 DPI, expression of several interferon stimulated genes peaked in addition to canonical inflammatory genes, notably *IRF8, IRF1, IRF7, IFI27,* and *IFI6*. This pattern of gene expression was evident in both infiltrating macrophage subsets (**Figure 6D**). In the proliferating AM and AM subset, additional genes associated with tissue repair, apoptosis, and regulatory function are also upregulated (e.g., *PYCARD, CASP1, CARD9, TREM2*) with infection (**Figure 6D**).

**Figure 6:**
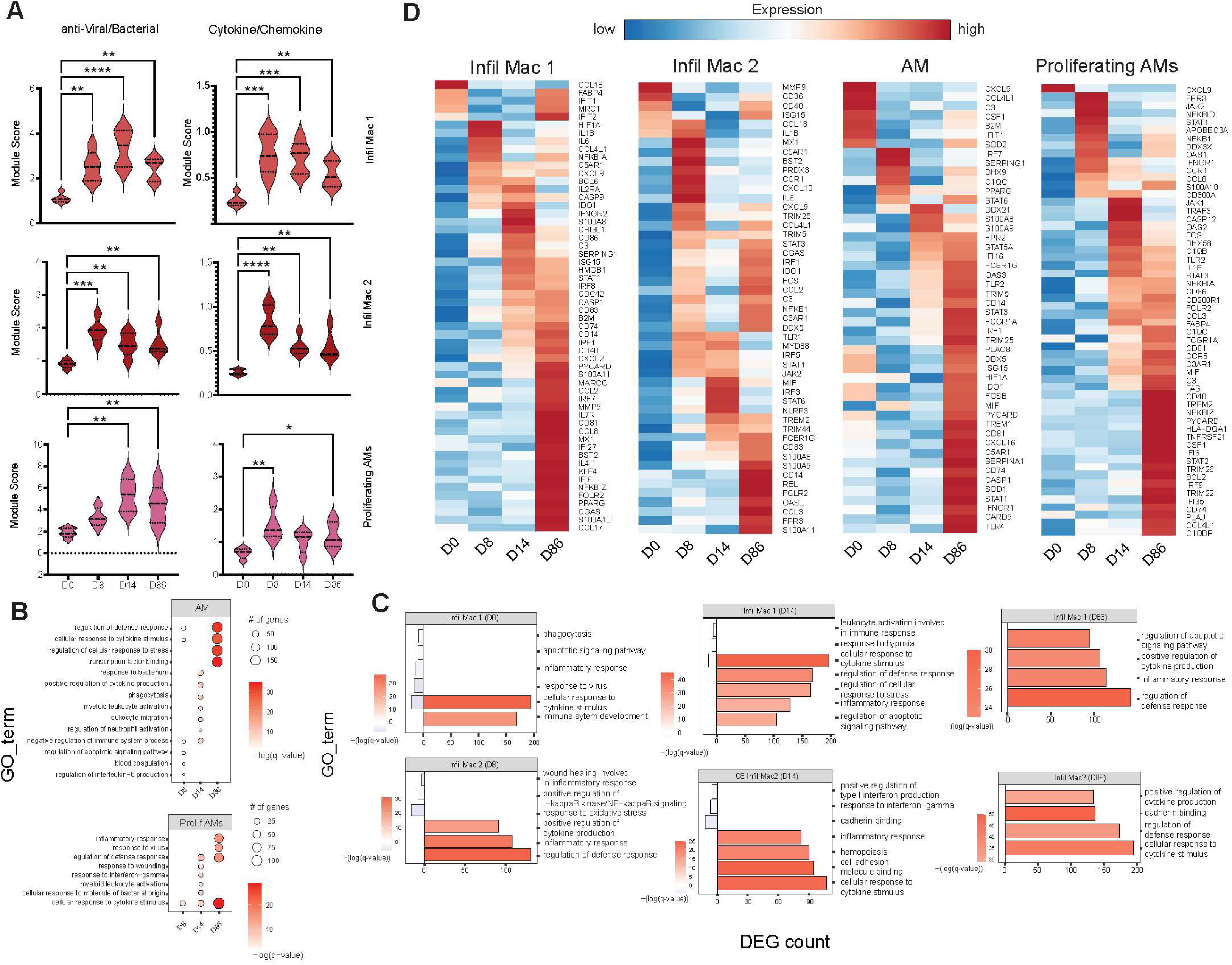
Single cell RNA sequencing of myeloid cells shows an association with defense against intracellular pathogen. **(A)** Violin plots showing the change in module scores of macrophage clusters across time. **(B)** Bubble plots of the gene ontology (GO) terms after functional enrichment of AM and proliferating AM clusters. The size of the bubble indicates the number of genes associated to a GO term and color indicates the significance compared to 0 DPI. **(C)** Bar graphs of the GO terms after functional enrichment of the infiltrating macrophage clusters at the indicated DPI. The magnitude of the bar indicates the number of genes associated with the GO term, color indicates the significance, and directionality indicates upregulation (positive) or downregulation (negative). **(D)** Heatmap of significant differentially expressed genes within macrophage clusters. Scaled average gene expression is represented by the color within each heatmap from low (blue) to high (red). Significance for panel A was calculated using a 1-way ANOVA with Benjamini and Hochberg’s post-hoc FDR comparisons relative to 0 DPI. *p < 0.05, **p < 0.01, ***p < 0.001, ****p < 0.0001.

To uncover transcriptional changes within the lymphoid clusters, we first measured changes in module scores associated with cytotoxicity of CD8 T cells, ISG T cells, and NK cells (**Figure 7A**). Cytotoxicity potential significantly increased at 8 and 14 DPI within ISG activated T cells and NK cells, DPI 14 for CD8 GNLY, and 8-86 DPI in ANKRD11+ T cells (**Figure 7A**). Functional enrichment of DEGs detected within CD8 ANKRD11+ subset showed over-representation of GO terms associated with antimicrobial responses at 8 DPI such as “response to virus” and “cellular response to cytokine stimulus” followed by a shift to “leukocyte migration” and “response to oxidative stress” 14 DPI (**Figure 7B**). Notable DEGs at 8 and 14 DPI include several ISGs (*STAT1, DDX60, TRIM22, MX1 and IFI27*) and genes in response to oxidative stress in the latter timepoint (*HIF1A, COX1, PROX6*) (**Figure 7C).** DEG within the CD8 GZM+ subset did not significantly enrich to GO terms and mostly consisted of ISGs (*STAT1, IRF1, DDX5)* and cytolytic molecules *(GNLY, KLRB1*). Within the S100A4+ CD4 T cell cluster, DEG mapped to GO terms ranging from “lymphocyte differentiation” and “response to virus” 8 DPI, “regulation of defense response” 14 DPI, and finally “transcription factor binding” indicative of maturation of the T cell response (**Figure 7B**). Notable DEG in this subset include ISG (*ISG15, ISG20, STAT3, CASP1, JAK1, IRF1)*, T cell signaling (*CD3G, CD3D, LCK, CD40LG)*, as well as genes important for type II IFN signaling *(IFNG, IFNGR1*) (**Figure 7C)**. We did not observe any notable DEGs in the other T cell and NK subsets.

**Figure 7:**
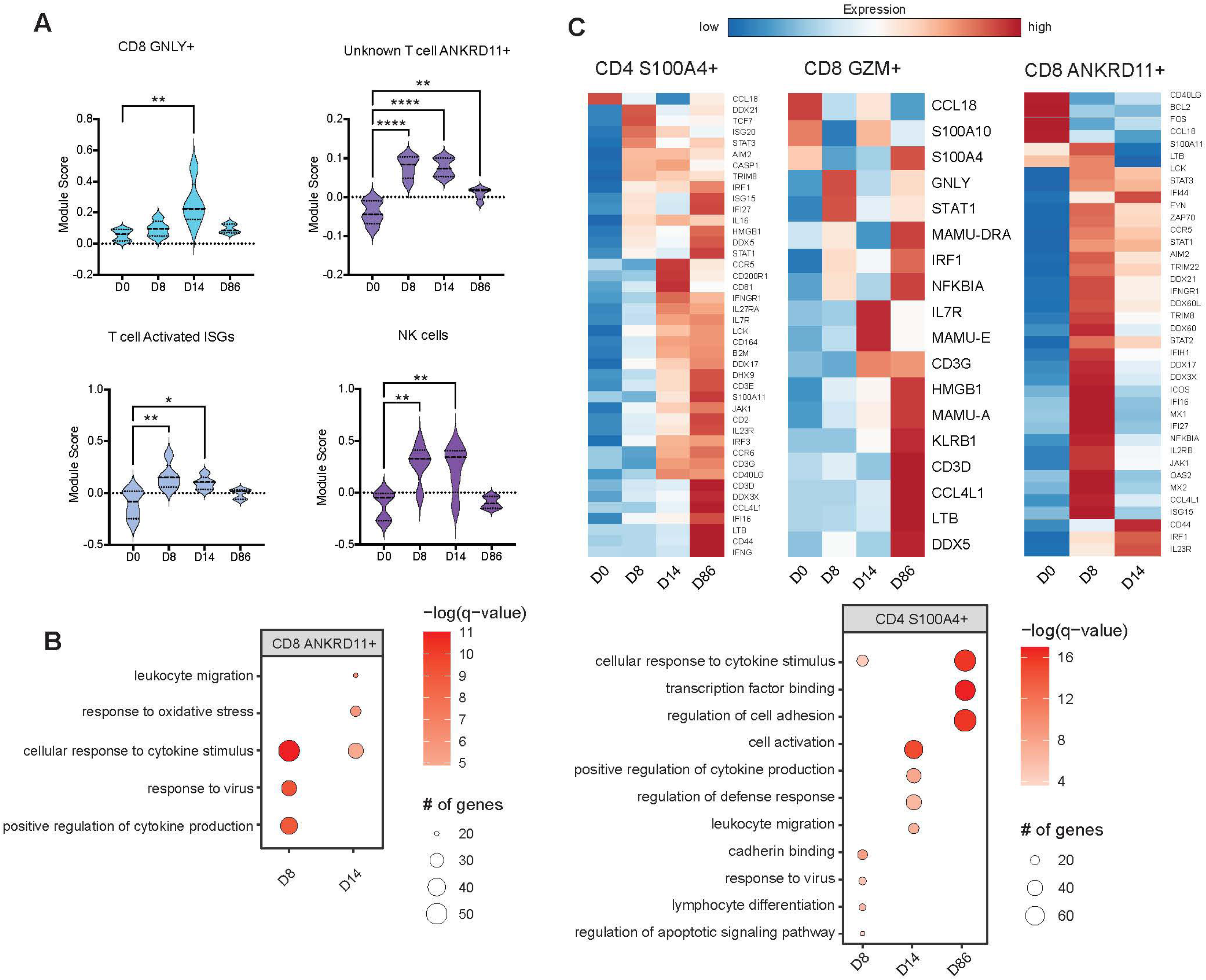
Single cell RNA sequencing of lymphoid cells indicates a CD4 and CD8 T cell response to MAC infection. **(A)** Violin plots showing the change in module scores associated with cytotoxicity of notable lymphoid clusters across time. **(B)** Bubble plots of the gene ontology (GO) terms after functional enrichment of CD8 ANKRD11+ and CD4 S100A4+ clusters. The size of the bubble indicates the number of genes associated to a GO term and color indicates the significance compared to 0 DPI. **(C)** Heatmap of significant differentially expressed genes within notable lymphoid clusters across time. Scaled average gene expression is represented by the color within each heatmap from low (blue) to high (red). Significance for panel A was calculated using a 1-way ANOVA with Benjamini and Hochberg’s post-hoc FDR comparisons relative to 0 DPI. *p < 0.05, **p < 0.01, ***p < 0.001, ****p < 0.0001.

We next compared the transcriptional profile of the young and aged cells within each cluster. A significant number DEGs were only detected within infiltrating and alveolar macrophages, CD4+S100A4+, and NK cells (**Supp Figure 7**). Functional enrichment of DEGs detected within infiltrating and alveolar macrophage clusters revealed processes important for inflammation, responses to intracellular pathogens, and apoptosis (**Supp. Figure 7A**). Some of the notable DEGs upregulated in the aged animals include ISGs (*IRF*s, *IFI*s, *OAS2*, and *DDX5*), transcription factors (*STAT1, NFKBIA*)*, CD74*, and *CASP1* (**Supp. Figure 7B)**. DEGs within the CD4 S100A4+ cluster mapped to TCR and IFN signaling (*CD3E, CD40LG, IRF1*) (**Supp. Figure 7A, C**) while DEGs within the NK subset enriched to cytolytic and antiviral pathways (*NKG7, KLRD1, KIR2DL4, GNLY*) (**Supp. Figure 7A, C**).

### MAC pulmonary disease results in the loss of compartmentalization of the lower respiratory microbiome

Prior studies reported significant shifts in the respiratory microbiome of individuals with NTMPD [15, 16]. Therefore, we profiled the right and left BAL microbiomes, as well as the nasal, oral, and gut microbiomes using 16S rRNA gene amplicon sequencing longitudinally. Plotting the percent relative abundance of bacterial phyla at all timepoints revealed striking differences between the fecal, nasal, oral, and lung microbial communities. The fecal community was dominated by Spirochaetota, Firmicutes, Campilobacterota, and Bacteroidota (**Supp. Figure 8A**). The respiratory communities did not have Spirochaetota or Campilobacterota present but was dominated instead by Proteobacteria. The nasal and lung communities harbored Actinobacteriota, which is the phylum containing *Tropheryma*, a prominent lung commensal in rhesus macaques (**Supp. Figure 8A**). Notable genera in the fecal microbiome are *Treponema*, *Lactobacillus*, *Methanobrevibacter*, *Prevotella*, and *Helicobacter* (**Supp. Figure 8B**). The nasal microbiome mainly consisted of *Dolosigranulum*, *Corynebacterium*, and *Psychrobacter* (**Supp. Figure 8B**). There were distinct dominant genera in the oral (*Rodentibacter, Gemella, Alloprevotella*) and lung (*Moraxella, Tropheryma*) microbial communities, but there were many overlapping genera such as *Porphyromonas*, *Neisseria*, *Fusobacterium*, and *Actinobacillus* (**Supp. Figure 8B**). Principle coordinate analysis (PCoA) shows that the fecal, oral, and nasal microbiomes are distinct while the BAL microbial community was intermingled with the nasal and oral microbiomes (**Supp. Figure 8C**). Following infection, the number of observed amplicon sequence variants (ASVs) was significantly increased in the right BAL samples at 8, 14, and 28 DPI (**Figure 8A**). Despite the lack of evidence of MAC spread to the left lung, a modest increase in the number of observed ASVs was also detected at 8 DPI (**Figure 8A**). On the other hand, the relative species diversity of the nasal, oral, and fecal microbiomes was unaffected by MAC infection (**Supp. Figure 8D**). Moreover, beta diversity was significantly decreased in the right BAL early in infection as evident by the diminished unifrac distance (a metric for inter-species relatedness) at 8 DPI (**Supp. Table 3**).

**Figure 8:**
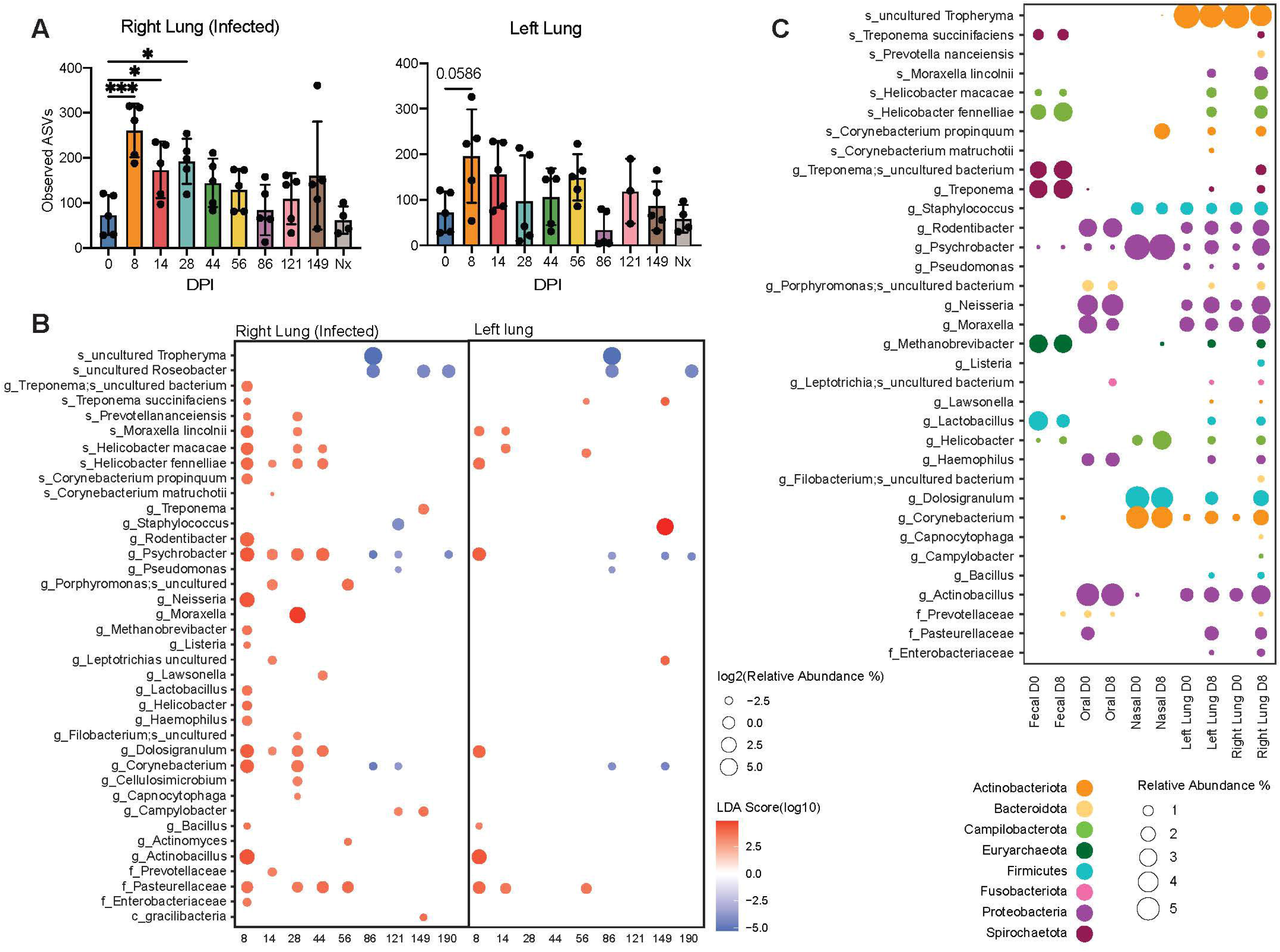
Microbiome profiling reveals microbial decompartmentalization of the gut and upper and lower respiratory tract. **(A)** Observed amplicon sequence variants (ASVs) in the right and left lung as the disease progresses. **(B)** Bubble plot derived from LEfSe analysis of the microbiome profiling of the right and left BAL. The size of each bubble represents the percent relative abundance and color represents the linear discriminant analysis (LDA) score. **(C)** Bubble plot of the most abundant species, family, or genera at each location at day 0 and day 8. The size of each bubble represents the percent relative abundance and color represents the Phyla.

To elucidate specific microbial shifts in left and right lung over the course of infection, Linear Discriminant Effect Size (LEfSe) analysis was carried out, which revealed an increase in the relative abundance of numerous bacterial species within the right lung microbiome between 8 and 56 DPI, notably several *Helicobacter* species (*H. macacae* and *H. fennelliae*) and *Methanobrevibacter*. Other species decreased in abundance 86-190 DPI, most notably, *Tropheryma* and *Roseobacter* (**Figure 8B**). Shifts in microbial communities were less dramatic in the left lung; nevertheless, increased abundance of *Actinobacillus, M. lincolnii, H. fennelliae* were noted 8 and 14 DPI (**Figure 8B**). Interestingly, several of the genera and species that increased in abundance over the initial 56 DPI are typically found in fecal (*Helicobacter*, *Treponema*, *Methanobrevibacter*, and *Lactobacillus*), oral (*Haemophilus*) and nasal (*Corynebacterium*, *Dolosigranulum*, and *Psychrobacter*) communities suggesting a loss of strict compartmentalization with MAC infection (**Figure 8C**). Specifically, at 8 DPI, members of Actinobacteriota (*Corynebacterium*), Proteobacteria (*Psychrobacter*, *Neisseria*, and *Moraxella lincolnii*), Campilobacterota (*Helicobacter*), Spirochaetota (*Treponema*), and Firmicutes (*Lactobacillus*, *Dolosigranulum*, and *Bacillus*) increased in abundance compared to baseline (**Figure 8C**).

## DISCUSSION

We used a rhesus macaque model to evaluate the pathophysiology of acute pulmonary MAC infection. This model closely recapitulates the hallmarks of human pulmonary MAC with a more severe phenotype in aged animals, which is exhibited via the radiologic findings, gross pathology, and immunohistochemistry. Our data highlighted important changes in immune responsiveness, with immune infiltrates flooding into both lungs despite the lack of immune mediators in the uninfected lung, as well as a prominent transcriptional shift towards combating intracellular infections. Microbiome profiling of the nasopharyngeal cavity, lung, and feces revealed dysbiosis of the lung microbiome immediately after infection. Indeed, differentially abundant bacterial DNA observed in the lungs was determined to be from species normally found in the oral, nasal, and gut microbiomes, indicating a loss of compartmentalization.

Despite the small group size, a more severe disease was noted in aged compared to young animals in the form of pulmonary consolidation and lesions, which is in line with established clinical literature [61–63]. Increased lung damage in aged animals was also evident from histological analyses that revealed presence of lymphoid aggregates, interstitial fibrosis, pulmonary edema, and bronchopneumonia. Interestingly, morphological and histological changes were comparable during acute infection, but their severity increased at much later time points in aged animals long after live bacteria were no longer detected in the BAL. This suggests that tissue injury could be induced by aberrant immune activation rather than by bacterial replication. Alternatively, it is possible that low level of bacterial replication could be occurring in the epithelial cells within a biofilm, as patients with a history of NTMPD are known to have relapses or re-infection [64–69]. Mycobacterial biofilms are also known to form at the air-media interface [70–73]; therefore, investigating the epithelial cells of the lung for the presence of MAC will be essential in future studies. In line with this, our data show that MAC bacterial burden decreased very rapidly in the infected right lung and did not spread to the left lung. Our inability to detect live organisms beyond 8 DPI could be due to the robust inflammatory response that developed 8 and 14 DPI that could have eliminated the bacteria. This is especially relevant for MAC, a slow growing mycobacterium.

MAC-specific IgG levels were detected 14 DPI in the plasma, while their levels in the BAL show a modest transient increase suggesting that IgG may be important to limit dissemination. The role of IgG may be limited within the lungs since surfactant proteins (SP) A and D may opsonize mycobacteria [74]. Other studies showed that NTM infections result in high levels of circulating IgG and IgA which can be used to diagnose NTM infection [75, 76]. Moreover, it has been shown that antibodies have a functional role in the control of TB [77]. Notably, the increase of plasma IgG appears to be transient, potentially due to strategies mycobacteria employ to evade and subvert antigen presentation on MHC class II molecules [78–81] and the ability of the obligate intracellular NTM to inhibit cross-presentation on MHC class I molecules [82].

While the small group size precluded us from carrying out extensive comparisons between the two age groups, some interesting trends emerged. In line with the lack of dissemination from the original site of infection to the left lung, significant changes in soluble immune mediators were only observed in the right side. Overall, these changes were indicative of a response to an infection with increased levels of chemokines to recruit T cells (CXCL11, CXCL10), eosinophils (CCL5), immature dendritic cells and B cells (CCL20). These changes correlated with increased recruitment of T cells into the alveolar space. We also detected increased levels of growth factors important for recruiting monocyte-derived macrophages (GM-CSF) that coincided with infiltration of macrophages into the lung. Moreover, levels of cytokines involved in activating macrophages (IFNγ), defending against extracellular (IL-17) and intracellular pathogens (IFNα), Th1 responses (IL-12), and endothelial activation (TNFα). Interestingly, a closer examination revealed that IL-10 and CD40L were modestly lower in aged animals. Decreased levels of IL-10 could result in sustained inflammatory responses leading to the increased lung damage in aged animals [83]. Lower levels of CD40L indicates reduced expression of co-stimulatory molecules [84]. Finally, growth factors that play a critical role in wound healing (VEGF, PDGF-BB) were elevated later in the disease process in line with the initiation of post-infectious repair phase.

Notably, despite the lack of significant increase in immune mediators in BAL supernatants of the left lung, immune cells infiltrate both lobes after infection. Flow cytometry and single cell RNA sequencing revealed similar patterns over the course of infection for both myeloid and lymphoid cells. Relative abundance of AMs precipitously declined due to the influx of T cells. While their levels do not recover over the course of 86 DPI, infiltrating macrophages are recruited into the alveolar space. We were also able to identify a cluster of proliferating AMs. This finding is in line with studies showing proliferation of AMs in mice and humans following significant loss [85–87]. Expression of genes important for anti-viral/bacterial responses, cytokine-chemokine signaling, and wound healing were upregulated significantly in all macrophage subsets throughout 86 DPI. Differential gene expression revealed heightened inflammatory signatures in macrophages from aged compared to young animals, including sustained high expression of several ISGs. On the other hand, expression of genes important for antigen presentation, co-stimulation of T cells, and type II IFN signaling were upregulated in young animals. Collectively, these data indicate a dysregulation of anti-microbial function with age in lung resident macrophages [88].

Relative abundance of natural killer (NK) significantly increased during the late stages of infection (121-190 DPI), long after the bacteria was no longer detected in the BAL. This influx of NK cells was evident. Following cell-to-cell contact, NK cells can activate MAC-infected macrophages to kill the pathogen via the secretion of TNFα [89] and IFNγ [90]. Moreover, neutralizing IFNγ exacerbates mycobacterial growth while inhibiting NK-mediated cytolysis has no effect on bacterial growth [91]. Interestingly, NK cells of young animals expressed higher levels of cytolytic genes such as *GNLY* and *GZMB* whereas NK cells in aged animals expressed higher levels of alarmins and MHC-I molecules.

A large influx of T cells, notably CD4 T cells, is detected early in the infection and their levels remained elevated throughout the course of infection in both lungs, a finding in accordance with the critical role of CD4 T cells in fighting mycobacteria [92, 93]. Interestingly, the frequency of MAC-specific CD4 T cells was higher in the left lung. This response was dominated by IFNγ and TNFα while IL-17 producing CD4 T cells were not consistently detected in either lung [88]. In contrast, MAC-specific CD8 T cell responses were larger in the right lung compared to the left lung and were more significant in the young animals compared to the aged. The larger CD8 T cell responses may be explained by the ability of live mycobacteria to inhibit MHC II antigen processing and presentation pathways resulting in cross presentation on MHC I [78–81, 94].

Single cell RNA sequencing revealed several T cell states. The ISGs T cell cluster peaked during the acute phase in line with the peak levels of IFNα detected in the BAL. ISGs are crucial for the host to combat intracellular pathogens including mycobacteria [95, 96]. Other T cell subsets that peaked during acute phase included CD8 expressing high levels of granulysin and ANKRD11. Granulysin is known to play a role in clearing intracellular bacteria, including mycobacteria [97, 98]. The CD8 ANKRD11+ T cell cluster expressed high levels of genes critical for lymphocyte activation, migration, T cell signaling, and histone modification. Finally, two different CD4 T cells clusters (CCR6+ and S100A4+) appeared later in infection in young and aged animals, respectively. The presence of CCR6+ CD4 T cells in young animals. CCR6+ CD4 T cells were recently reported to be important for prevention of Tb-associated immune reconstitution syndrome and their detected in young animals only is aligned with better disease resolution in these animals [99]. On the other hand, the presence of CD4+ t cells expressing alarmins in aged animals suggests a dysregulated inflammatory response.

Significant disruption in the BAL microbiome was noted in both lungs that were sustained beyond 8 DPI (last time point at which live bacteria were detected). Disease states such as cystic fibrosis or bronchiectasis often cause dysregulation of the normal lung microbiota which is characterized by increased microbial biomass, but lower diversity [16, 100]. In contrast, our data show that MAC infection leads to increased diversity in the microbial community of the lungs and is accompanied by detection of several species normally sequestered in other anatomical sites, including the gut. This finding correlates to clinical NTM disease where a larger proportion of anaerobic bacteria, such as *Prevotella* and *Fusobacterium*, can be found [101]. Notable species that were determined to be differentially abundant via 16S rRNA sequencing include the enrichment of microbes predominantly found in the nasopharyngeal (*Dolosogranulum, Corynebacterium*) and oral (*Neisseria, Moraxella*) tracts as well as the fecal microbiome (*Helicobacter fenneliae, Treponema, Methanobrevibacter*). Additionally, a depletion of *Tropheryma* at 121 DPI was observed. Our group has previously shown that *Tropheryma* is a major component of a healthy rhesus macaque lung microbiome [48]. This dysbiosis suggests that the inflammatory changes induced by MAC infection may have led to loss of compartmentalization. However, it remains unclear whether these data translate into dysbiosis of live organisms or of bacterial DNA fragments that can now access the lung due to the impaired barrier (driven by inflammation). Regardless, bacterial products can activate myeloid cells, thereby exacerbating inflammatory responses and further damaging tissues [102, 103]. This could explain the heightened activation of AM and infiltrating macrophages 3 months after infection.

In summary, data presented in this study show that rhesus macaques recapitulate the hallmarks of NTMPD including bronchiectasis and increased disease susceptibility with age. Our observations suggest that the response to MAC is mediated by both macrophages and T cells. Interestingly, some of the lung tissue damage may be mediated by the immune response rather than the bacterium. While the availability of this model will facilitate future studies that interrogate host-pathogen interactions as well as development and testing of novel therapeutics, it is not without limitations. The small cohort size precluded an in-depth comparison of the immune response in young versus aged animals. This study focused mainly on the immune response in the lung. Additionally, a single bolus of MAC in only the lower lobe of the right lung was deployed in this study. Therefore, future studies will use a smaller inoculum or a different method of instilling MAC such as an inhalation chamber to deliver the bacteria into a larger area of the lung in order to more closely resemble human exposure. In addition, this approach would potentially reduce the magnitude of the initial inflammatory response, thus allowing the bacteria more time to establish infection. Future studies will also use larger cohorts to allow comparison age and sex-comparisons as well as evaluation of systemic responses.

## Figure Legends

**Supp. Figure 1: Electron Microscopy reveals *M. avium* residing in biofilms and macrophage. (A)** SEM showing *M.avium* (PCR confirmed) embedded in left lung biofilm. Arrows showing bacteria. **(B)** SEM showing *M.avium* (PCR confirmed) embedded in matrix rich biofilm in the right lung. Arrows showing bacteria. **(C)** SEM showing Plug in the right lung showing tick secretion and many host cells. **(D)** TEM showing a lung macrophage in the airway plug with several viable *M.avium* inside of cytoplasm vacuoles. Arrows showing bacteria.

**Supp. Figure 2: Hematoxylin and eosin (H&E) stains of NHP lung samples.** Representative H&E images at the indicated magnification of lungs from young and aged rhesus macaques infected with 6.8×10^8^ CFU of MAC. Arrows point out lymphoid aggregates.

**Supp. Figure 3: Immunofluorescent assay (IFA) of NHP lung samples.** Representative IFA images at 20x magnification of lungs from young and aged rhesus macaques infected with 6.8×10^8^ CFU of MAC. The fluorescent channels used are 4’,6-diamidino-2-phenylindole (DAPI) stain for a general DNA stain (Blue); enhanced green fluorescent protein (EGFP) conjugated to anti-CD3 for T cell identification (Green); Cyanine 5 (Cy5) conjugated to anti-CD20 for B cell identification (Yellow); Cyanine 3 (Cy3) conjugated to anti-CD68/163 for identification of macrophages and monocytes (Red).

**Supp. Figure 4: Immunofluorescent assay (IFA) segmentation algorithm allows for quantifying immune cells. (A-E)** The fluorescent channels for **(A)** 4’,6-diamidino-2-phenylindole (DAPI) stain; **(B)** enhanced green fluorescent protein (EGFP) conjugated to anti-CD3 for T cell identification; **(C)** Cyanine 5 (Cy5) conjugated to anti-CD20 for B cell identification; **(D)** Cyanine 3 (Cy3) conjugated to anti-CD68/163 for identification of macrophages and monocyteI**(E)** Merged fluorescent channels. **(F)** Example images of a single selected field of lung tissue followed by **(G)** the segmentation strategy from the HALO software algorithm used to quantify immune cells. **(H)** The average percent abundance of macrophages, T cells, or B cells found in the following areas of the lung grouped by young and aged: right middle, right cranial, right caudal, right accessory, left middle, left cranial, and left caudal lobes. Student’s T tests were performed between the young and aged groups. *p < 0.05, **p < 0.01, ***p < 0.001, ****p < 0.0001.

**Supp Figure 5: Representative gating strategy for flow cytometry analysis** Three experiments each with their own unique panel were ran at each timepoint to analyze the **(A)** innate and **(B)** adaptive immune systems and **(C)** intracellular cytokines after stimulation with 1 mg/mL MAC lysate. **(A)** The innate immune system was gated by initially viewing live, single cell lymphocytes followed by differentiating AMs and non-AMs. The CD3-CD20-population of the non-AMs were further delineated into monocytes, dendritic cells (DCs), and NK cells. **(B)** The adaptive immune system was analyzed by differentiating the live, single cell lymphocytes into B cells (memory, marginal zone, and naïve) and CD4 and CD8 T cells. The T cell subsets were further delineated into naïve, central, and effector memory T cells. The transcription factor Ki67 was used as a proliferative marker for each. **(C)** The intracellular cytokine staining assay was gated by differentiating single cell lymphocytes into IFNγ+TNFα+ CD4 and CD8 T cells. The IL-17+ CD4 T cell population was delineated. Additionally, the CD20-single cell lymphocytes were separated into monocytes and DCs.

**Supp Figure 6: Single cell RNA sequencing by timepoint and age g. (A)** UMAP of immune cells showing that each cluster from Figure 5 is equally represented by each sample. **(B)** Heatmap of marker genes for each UMAP cluster. **(C-E)** Bubble plots of notable GO terms from functional enrichment across all myeloid and lymphoid cell types. The size of the bubble indicates the number of genes associated to a GO term and color indicates the significance compared to 0 DPI.

**Supp Figure 7: Bubble plots from single cell RNA sequencing grouped by. (A)** Bubble plot of the GO term enrichment by age for the indicated cell subsets. The size of the bubble indicates the number of genes associated to a GO term and color indicates the significance compared between young and aged groups. **(B)** Bubble plots of differentially expressed genes (DEGs) for the infiltrating macrophage and AM clusters, or **(C)** for the CD4+S100A4+ T cell and NK cell subsets grouped by age category. The size of the bubble indicates the percent of cell expressing the indicated genes while color indicates an upregulation (red) or downregulation (blue).

**Supp Figure 8: Microbiome profiling of the BAL samples and nasal, oral, and fecal s. (A)** Phyla bar plot displaying the dominant phyla and their percent relative abundances at all timepoints at the indicated sites grouped by age category. **(B)** Bubble plot showing the dominant genera at all timepoints at the indicated sites grouped by age category. Size of the bubble indicates the number of genes associated with the corresponding genera. **(C)** The weighted and unweighted principal coordinate analysis (PCoA) plot of the BAL (0 DPI, left, and right), oral, nasal, and fecal swabs at the indicated DPI. The BAL samples at 0 DPI contained cells from both the left and right lung. **(D)** Observed ASVs of the nasal, oral, and fecal swabs at all DPI.

**Supp. Table 1: Metadata of the research animals investigated during this study**

**Supp. Table 2: Module score categories and the list of genes associated with each category**

**Supp. Table 3: The weighted intragroup Unifrac distances showing β-diversity at each time point and comparing to baseline (0 DPI) for the right and left BAL**

## AUTHOR CONTRIBUTIONS

N.S.R., S.G.K., E.R.S, K.W. and I.M. conceived and designed the experiments. M.D., S.G.K., and E.R.S. identified suitable animals and collected samples. N.S.R., I.R.C. M.D., E.G.N., C,F., and L.B. performed the experiments. N.S.R, I.R.C., E.G.N., and D.B.A. analyzed the data. I.R.C., E.G.N. and I.M. interpreted the results and wrote the paper. All authors have read and approved the final draft of the manuscript.

## ACKNOWLEDGMENTS

This work was supported, in part, by the intramural program of the National Institute on Aging, NIH (R01AI152258-04). This research was supported by the Biospecimen Procurement & Translational Pathology Shared Resource Facility of the University of Kentucky Markey Cancer Center (P30CA177558). The authors thank the ONPRC Division of Comparative Medicine and veterinary staff for their expert animal care. The authors also thank the veterinary technicians at the NIA nonhuman primate core for sample collection and support from the Division of Veterinary Resources animal care team (P51-OD011092).

