## Supplementary figures and images for "Impact of aging on the immunological and microbial landscape of the lung during non-tuberculous mycobacterial infection"

### Supplemental Figure 1

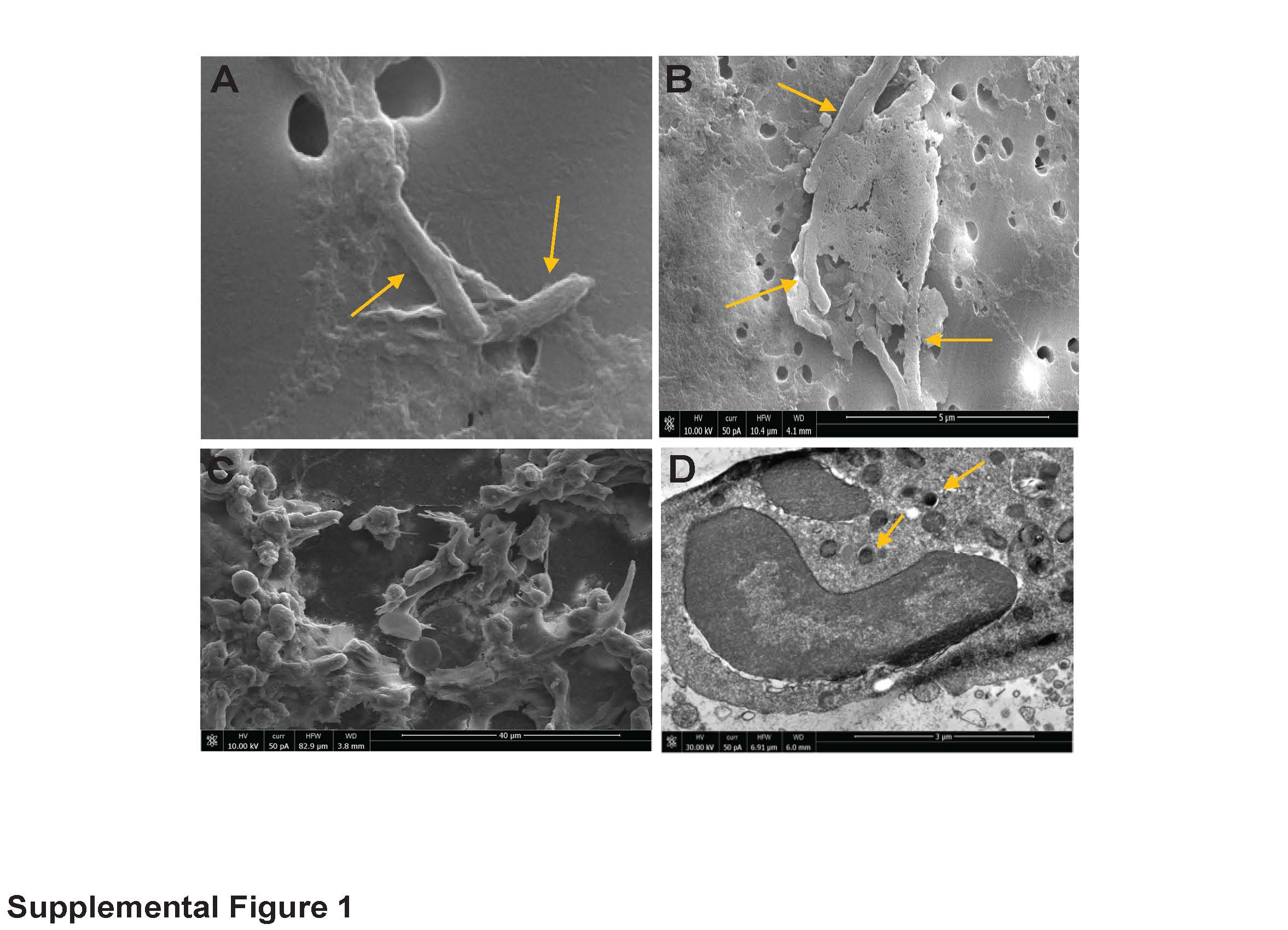

### Supplemental Figure 2

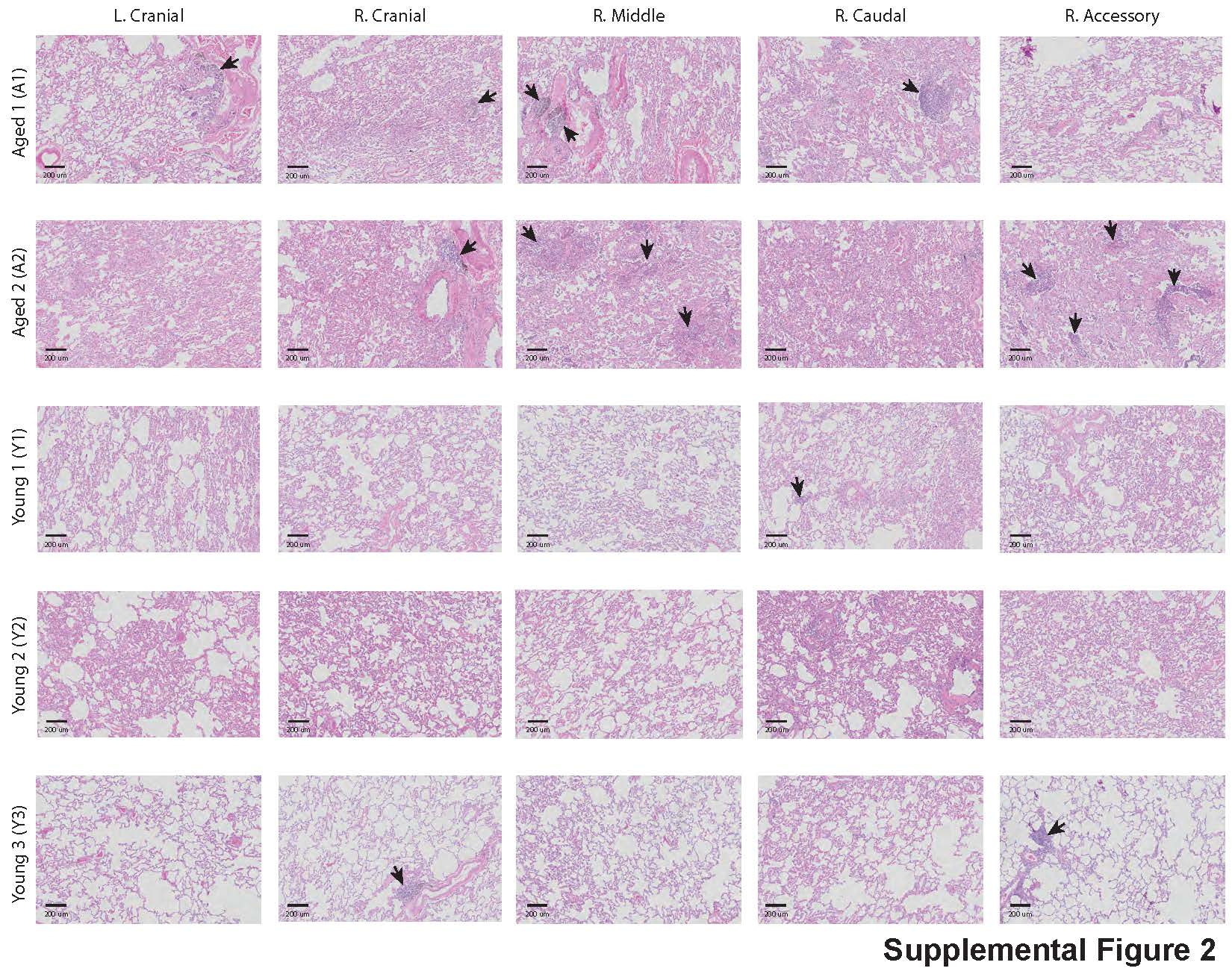

### Supplemental Figure 3

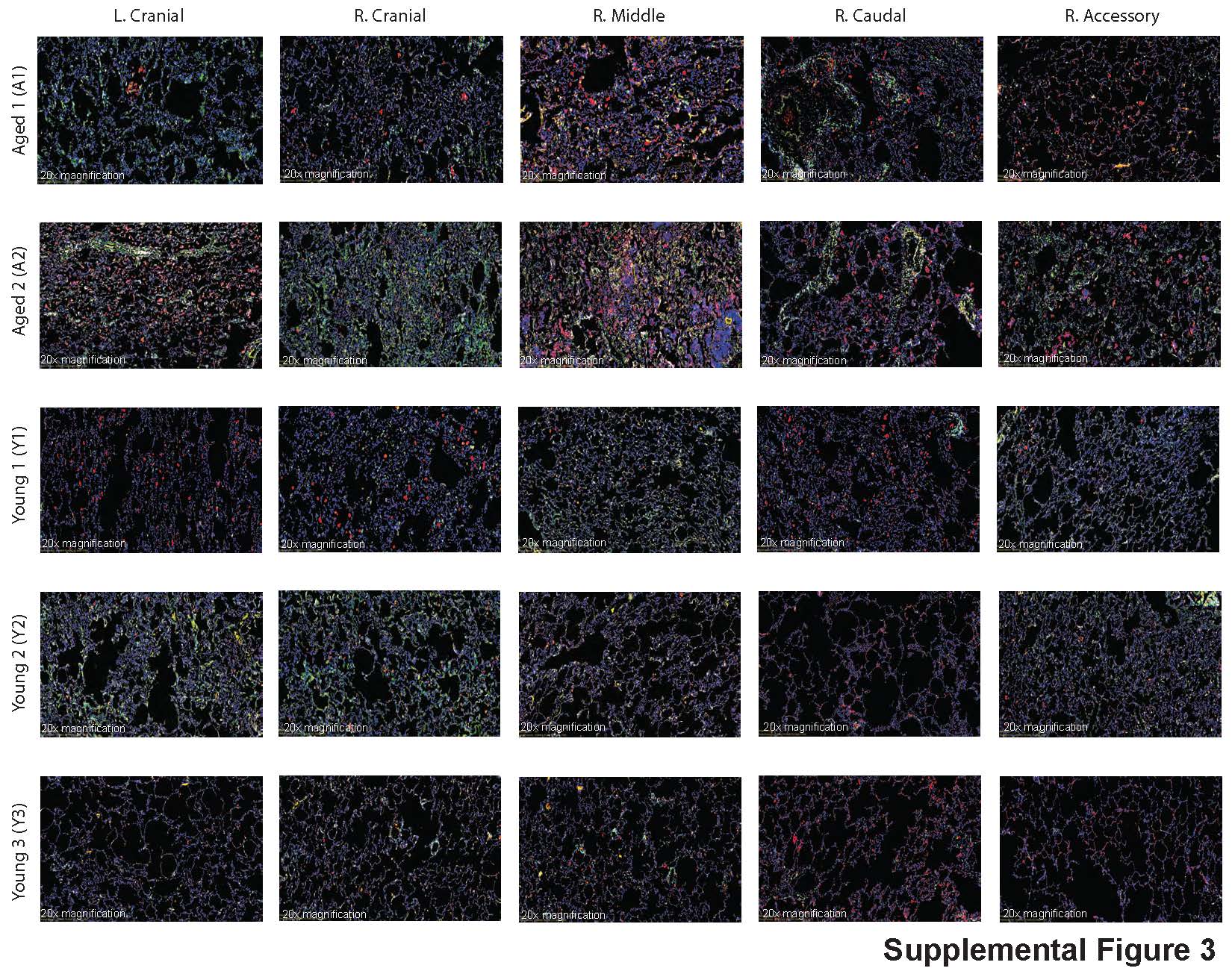

### Supplemental Figure 4

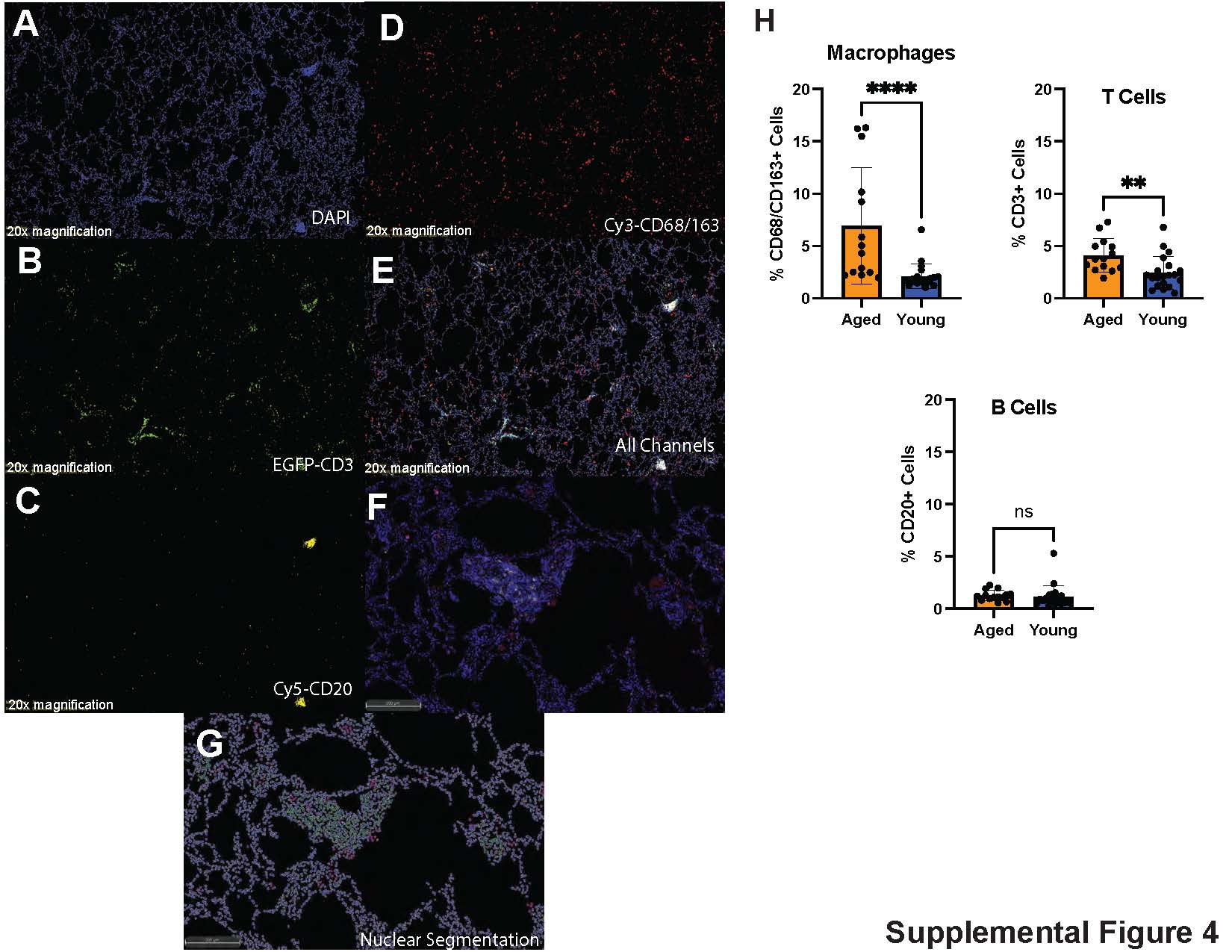

### Supplemental Figure 5

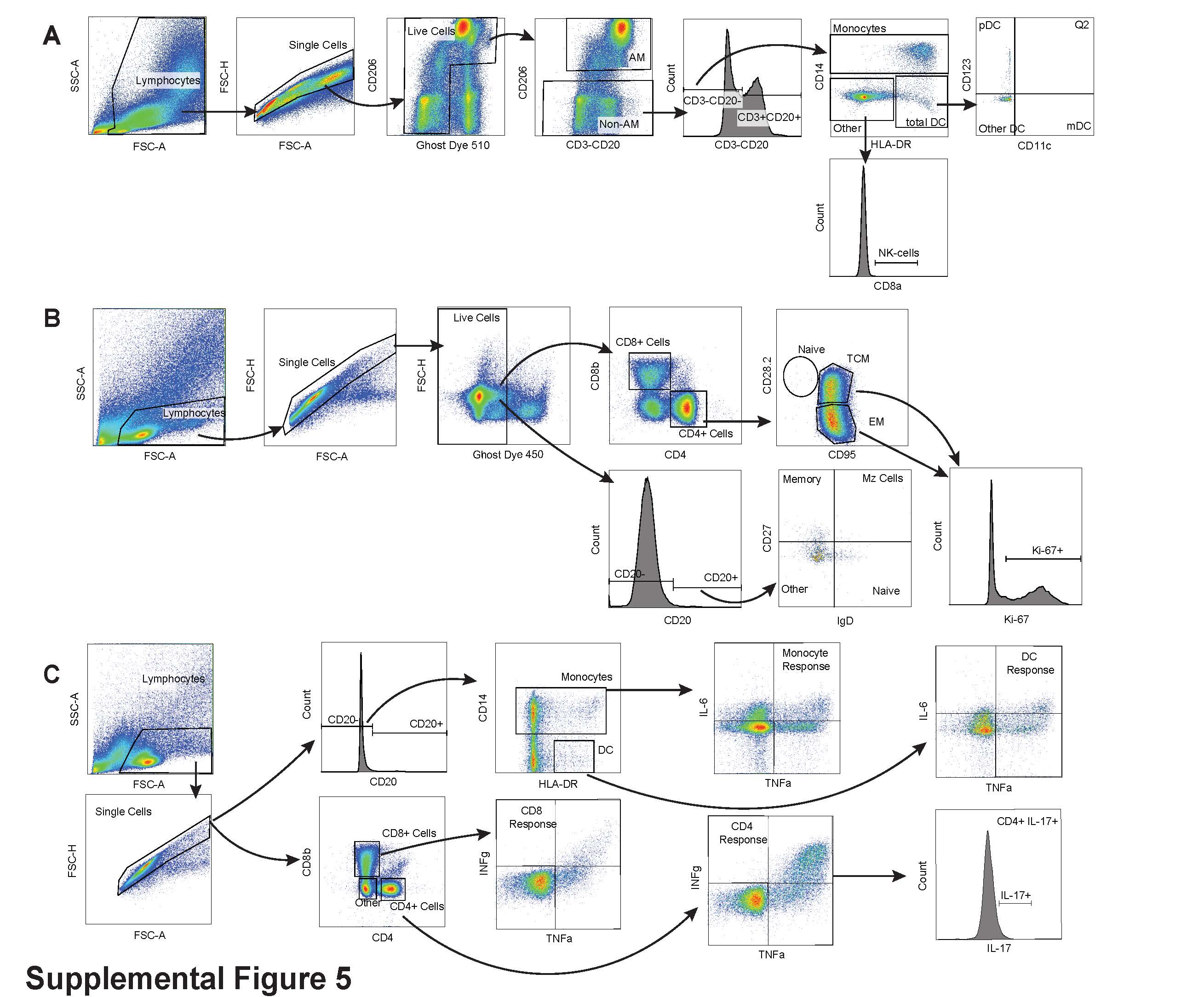

### Supplemental Figure 6

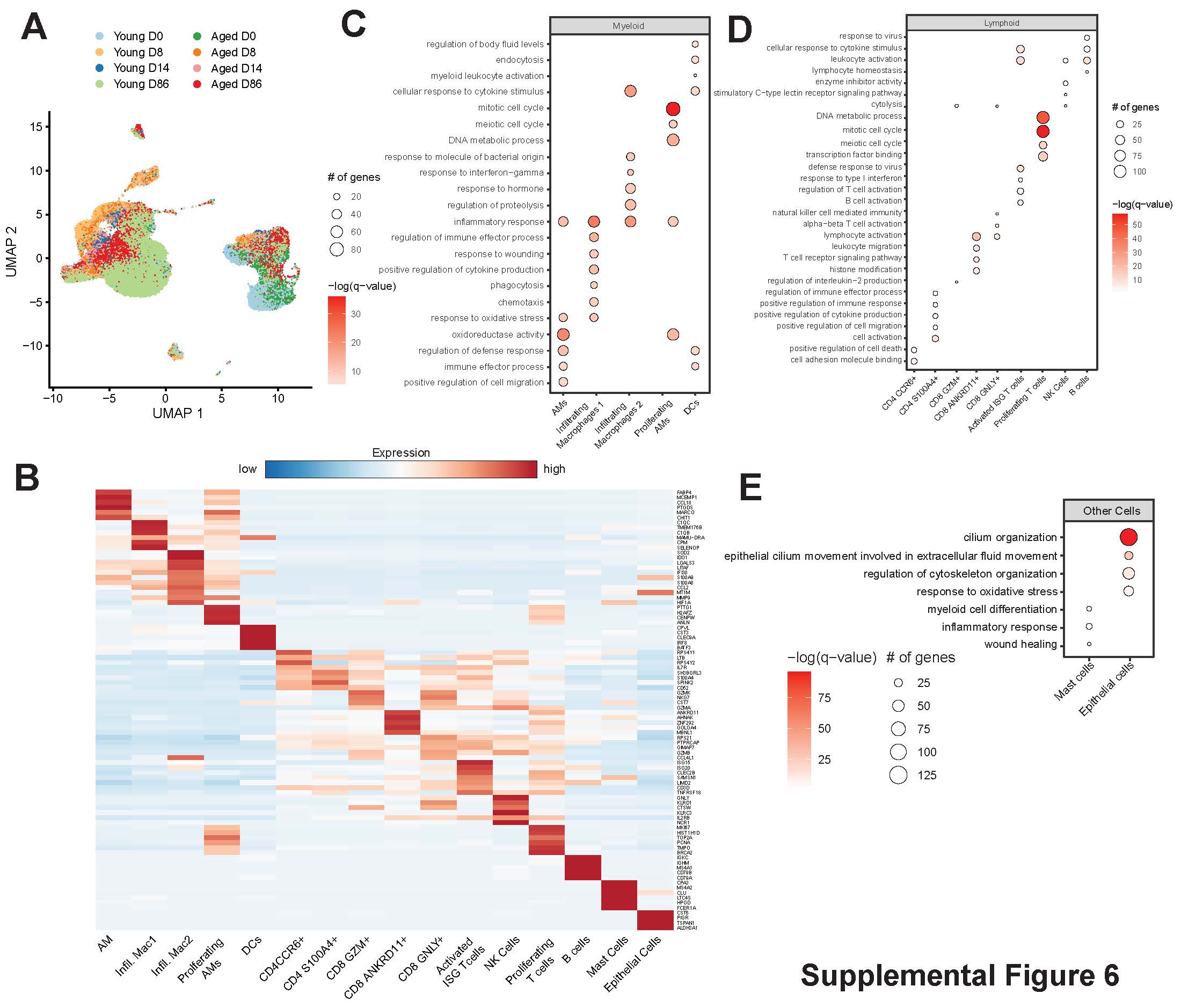

### Supplemental Figure 7

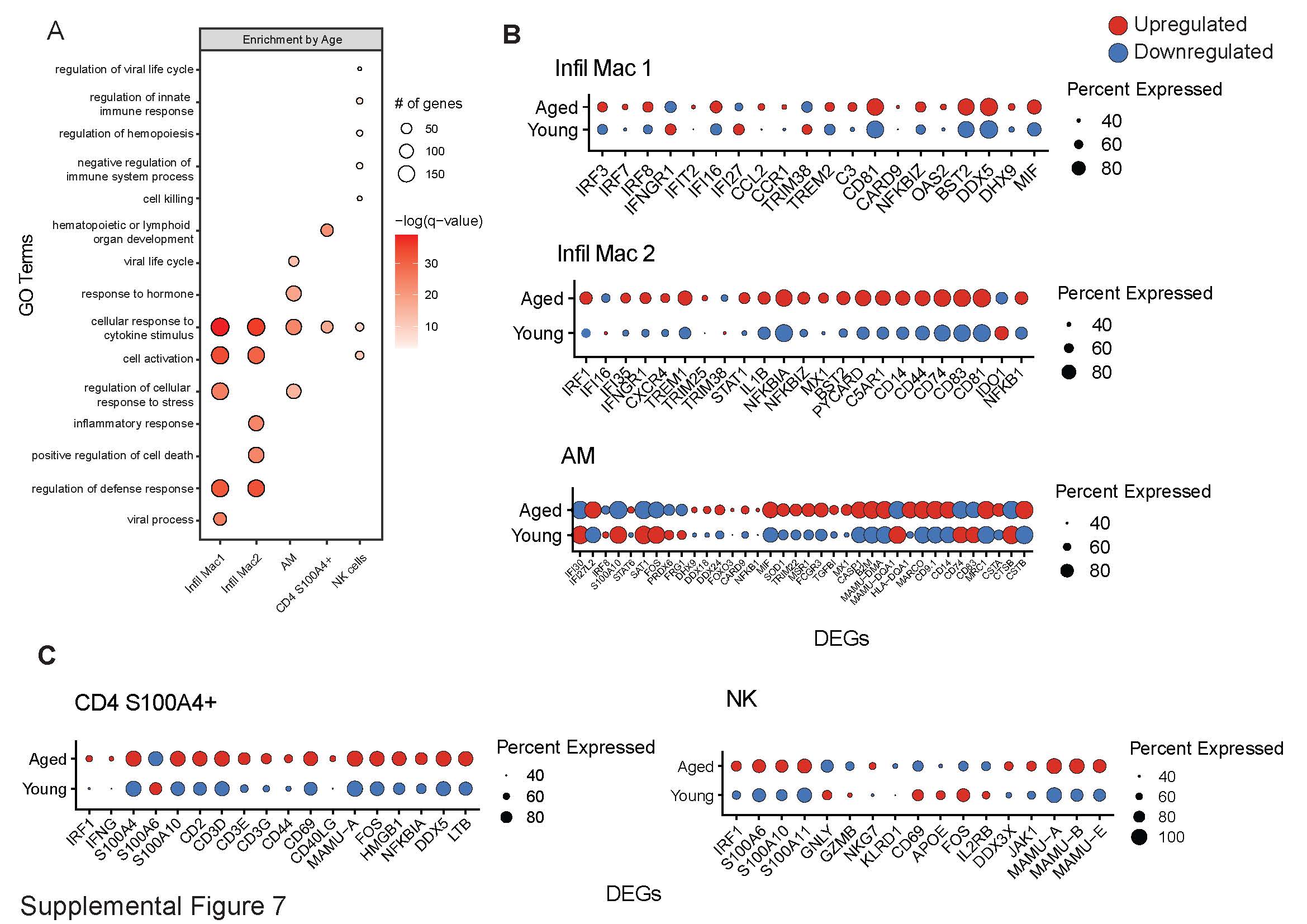

### Supplemental Figure 8

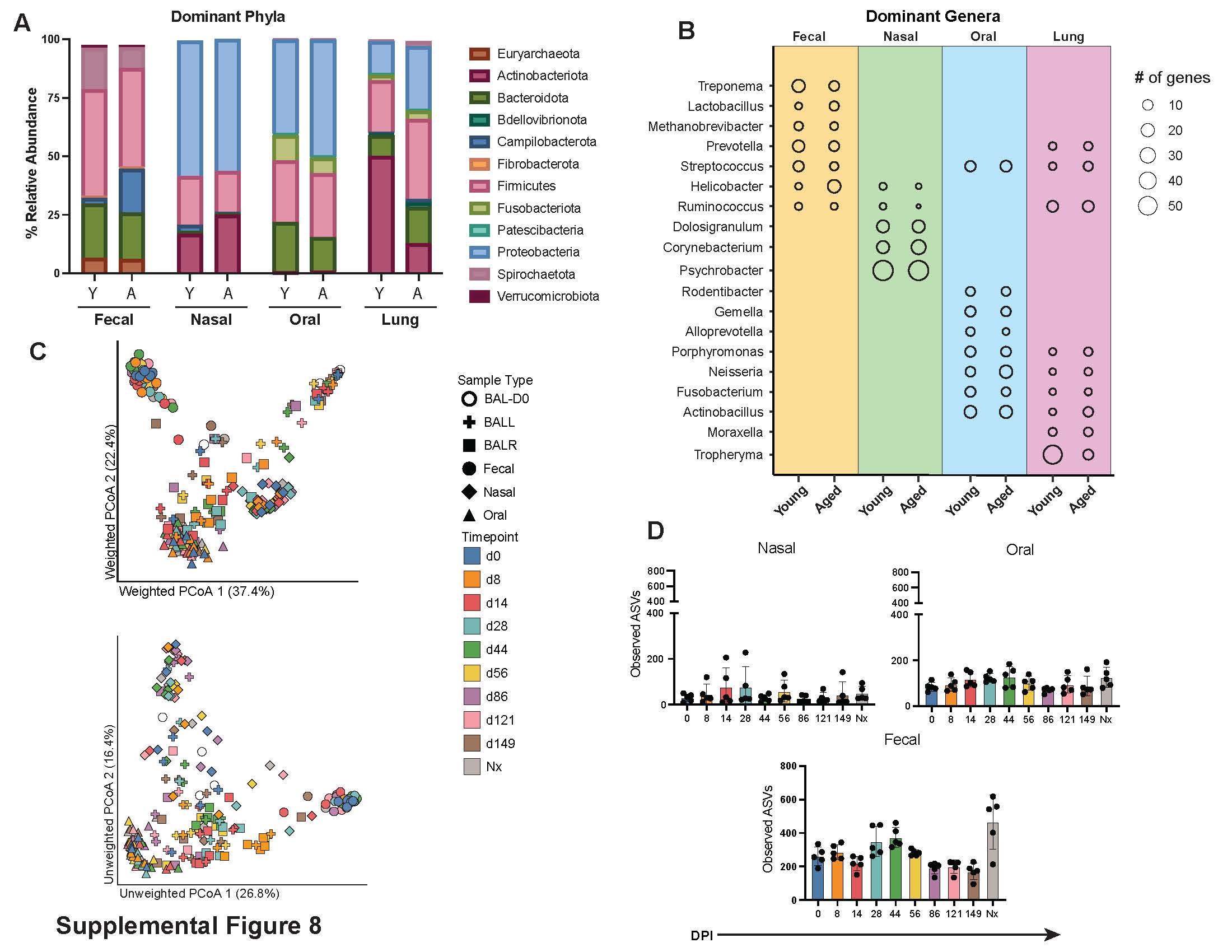
