## Supplemental Table 1 for "Impact of aging on the immunological and microbial landscape of the lung during non-tuberculous mycobacterial infection"

**Supp. Table 1: Metadata of the research animals investigated during thi**

| BlindID | Group | Age (y) | M/F | Weight at Inoculation (kg) |
|---|---|---|---|---|
| OR_Adult_3 | Adult | 8.82739726 | F | 8.70 |
| OR_Adult_8 | Adult | 6.0739726 | M | 7.10 |
| OR_Adult_10 | Adult | 5.69041096 | M | 8.00 |
| OR_Aged_12 | Aged | 21.0383562 | F | 7.80 |
| OR_Aged_32 | Aged | 23.9890411 | F | 8.65 |

**s study**

| Weight at Nx (kg) |
| --- |
| 8.45 |
| 7.45 |
| 7.10 |
| 8.05 |
| 9.10 |
