## Supplemental Table 2 for "Impact of aging on the immunological and microbial landscape of the lung during non-tuberculous mycobacterial infection"

**Supp. Table 2: Module score categories and the list of genes ass**

| anti_viral_bacterial_module_score_en | cytotoxicity_module_score_ensem |
|---|---|
| LTF | FCGR3 |
| TF | PTPRC |
| CAMP | IL7R |
| JCHAIN | KLRD1 |
| DEFB1 | PPP3CB |
| DEFA1B | CD1D |
| DEFA4 | STX7 |
| IGHM | CD1E |
| HIST1H2BJ | MAMU-DOB |
| RNASE6 | MAMU-DRA |
| RNASE7 | B2M |
| CTSG | IL12A |
| HLA-E | HLA-E |
| PLA2G6 | CYRIB |
| SEMG2 | IL12B |
| SLPI | IL23A |
| PLA2G1B | FADD |
| FAU | MR1 |
| BPIFA1 | SLC22A13 |
| AHSG | AGER |
| ITIH4 | NECTIN2 |
| INS | CTSH |
| FN1 | CTSC |
| LBP | EMP2 |
| HP | HPRT1 |
| SCARB1 | PRF1 |
| NECTIN2 | GZMM |
| ICAM1 | JAGN1 |
| SENP7 | GZMB |
| MX1 | SRGN |
| PHB2 | GZMA |
| RNF135 | STXBP2 |
| CLPB | LYST |
| OAS1 | RNF19B |
| ZDHHC1 | CD2 |
| PHB | PLEKHM2 |
| AKAP1 | KIF5B |
| NLRP1 | ULBP3 |
| IFIT1 | PTPN6 |
| CXCL10 | NKG2D |
| PGLYRP3 | PRDX1 |
| PGLYRP2 | IL18 |
| GPX1 | CEBPG |

DDB1
PLG
ELANE
GFI1
ZC3H12A
TNFRSF1B
CCR5
CD86
BCL10
PDE4B
WNT5A
CD80
PLSCR4
TNF
NFKBIL1
TNFAIP3
SERPINE1
CDC73
IL6
RHOA
TNFSF4
RIPK2
VIM
MRC1
CD36
HMGB2
XBP1
MAPK8
TRAF6
GIT1
JAK2
TNIP1
PDCD4
PLAA
NR1D1
ABCA1
CMPK2
IL10
RARA
PLSCR3
NR1H3
NR1H4
NKG2D
STAP1
CTR9
IL13
IL4
ARRB2
CLEC12B
HAVCR2
CD5L
SPI1
STAP1
CD160
SLAMF6
NCR3
IL21
LAG3
SH2D1A
RASGRP1
CD226
AP1G1
CRTAM
STAT5B
IL18RAP
CADM1
VAV1
SERPINB9
CR1L
LEP
PIK3R6
DNASE1L3
DNASE1

mml-mir-224
TRIM41
NOD2
LY96
IL12B
NUGGC
TICAM1
NFKBIB
NFKB1
MAPK14
AXL
DAB2IP
ADAM9
mml-mir-342
PAF1
IRF8
MALT1
ZFP36
PDCD1LG2
CDK4
NOS2
RELA
SBNO2
mml-mir-431
mml-mir-433
HAVCR2
CHMP5
IRF3
TNIP3
NLRP7
PYCARD
CD180
TLR4
LDOC1
CACTIN
ANKRD1
IL37
CD14
UPF1
ACOD1
CXCL1
CXCL6
CXCL8
AKAP8
ABL1

SIGLEC1
HRG
CFHR5
F2
REG3G
TPCN2
TPCN1
CTSL
HYAL2
GAS6
SFTPD
FGB
DYNLT1
GAPDH
PF4
NUCKS1
SAP30
SAP30BP
IFI27
FBXL2
TMEM41B
FMR1
PSMC3
RAB9A
APCS
PTX3
PRKN
ZBED1
CCNK
CCL8
VAPA
TRIM11
TRIM26
TRIM15
PML
TRIM21
TRIM28
TRIM35
TRIM8
TRIM13
MID2
TRIM32
MARCHF2
HDAC1
TARDBP

ZNF639
JUN
POU2F3
TFAP4
INPP5K
REST
HMGA2
SPINK5
ITGAV
CH25H
TRIM10
FCN3
TRIM59
SNX3
GSN
FCN1
MYD88
SCNN1B
AZU1
TUSC2
NLRP6
DHCR24
PSMB8
MAMU-DRA
MAML1
YTHDC2
TBC1D20
PPIB
ZNF502
CAV2
CSF1R
IGF2R
PC
CFL1
APOE
TAF11
RRP1B
SNW1
LEF1
CHD1
HPN
SMARCB1
CDK9
CCNT1
CTDP1

EP300
SP1
SMARCA4
PGC
KLK3
KLK5
KLK8
SYK
PRF1
CLEC7A
IFNG
FCER2
F2RL1
TMPRSS2
CD74
TRIM38
CD4
TMPRSS4
P4HB
EPS15
PIKFYVE
LRRC15
ACE2
FUCA2
EXOC2
EXOC7
HS3ST5
TRIM22
TRIM6
TRIM68
TRIM14
S100A14
S100A8
DUSP10
CD96
IL12RB2
IL23R
PTAFR
SNCA
LIAS
NOCT
ADH5
NFKBIA
PTGFR
MAPKAPK3

PRDX3
TRIB1
IL12A
TAB2
PELI1
ADAM17
TIRAP
CEBPB
MAPKAPK2
CPS1
SMAD6
PTGIR
CCR7
FER
CD6
F2R
RPS6KA3
MAPK1
SLC11A1
KCNJ8
GCH1
ERBIN
SRR
IDO1
CYP27B1
NOS1
IRAK1
PLCG2
UMOD
IRAK3
TNFRSF11A
PTGER1
TBXA2R
PTGER4
IL1B
OTUD5
MAPK3
WDR83
CHMP2B
CHMP7
CHMP4C
CHMP2A
CHMP4B
CHMP6
VPS4B

XPR1
AGTR1
CLDN1
GPR15
CLEC5A
NECTIN4
SIVA1
DAG1
CTSB
TYRO3
NRP1
AVPR1B
VAMP8
NPC1
GRK2
SMPD1
CLDN6
INSR
SLC3A2
UVRAG
AVP
CLEC4G
CD209
VPS18
DPP4
ARL8B
DDX39B
THOC7
THOC5
THOC3
THOC6
THOC1
THOC2
RAB7A
VPS37B
IST1
TSG101
HSP90AB1
NECTIN1
CD81

**sociated with each category**

cytokines_chemokines_module_sco
ZC3H12A
APOD
CHID1
PDCD4
PLA2G10
ADCY7
F2
ABCD1
MACIR
MEFV
APPL2
IL1R2
NLRP7
PPARA
PYCARD
KPNA6
APPL1
MYD88
TNF
IL6
IL17F
IL17A
GPSM3
PLA2G3
ANKRD42
HIF1A
mml-mir-324
CD6
IL17RA
CLEC7A
NOD2
TICAM1
IL17RC
CARD9
STAT3
TLR4
IL17B
BAP1
PLD4
ALOX5
PLD3
MAPK14
NOS2

LEP
HFE
SMAD7
FOXP3
ARG1
TBX21
IFNB1
MAP3K7
B2M
TRAF6
SASH3
FZD5
MALT1
TRAF2
IL18
IL1R1
IL18R1
IL1B
PRKCZ
DENND1B
GATA3
RSAD2
IL4
CD81
CCR2
TRPM4
CCR9
CCR1
CCR3
CCR5
CCR6
CCR4
CCR8
ACKR2
CCR7
ACKR3
GPR75
CCR10
CXCR5
CXCR3
CXCR6
CXCR2
CXCR1
GPR35
CXCR4

CX3CR1
CCRL2
CCL26
CCL19
CCL7
CCL16
CCL23
CCL18
CCL20
CCL17
MSMP
CCL27
CCL11
NARS1
CNIH4
DEFB1
CCL21
E2F8
OXSR1
RHOA
FOXC1
DUSP1
SLC12A2
RIPOR2
DOCK8
LOX
WNK1
SH2B3
LRCH1
STK39
CRNN
LDLRAP1
MME
HCLS1
GBP7
GBP3
HAX1
TRAF3IP2
POU4F2
CSF1R
CSF3
IL13
FOXF1
LEF1
TCIRG1

POU4F1
NFAT5
FOXH1
DPYSL3
CDK9
PID1
TPR
PYHIN1
RO60
PDE12
AXL
IFIT2
IFIT3
GAS6
UBE2G2
AIM2
IFI16
UBE2K
PNPT1
TRIM6
IRF1
HTRA2
OAS1
STAT1
STING1
CDC34
CDC42
EPRS1
CD58
STXBP3
VAMP3
WNT5A
ZYX
AIF1
SLC26A6
VIM
MRC1
KIF5B
TLR2
GAPDH
RAB7B
DAPK1
MYO1C
CDC42EP4
IL12B

VPS26B
STXBP4
RPS6KB1
GSN
IRF8
IL12RB1
AQP4
STX4
RAB20
WAS
STXBP1
STX8
ACTR3
ACTG1
ACTR2
RC3H1
HYAL3
HYAL1
HYAL2
MAPK11
HAS2
NKX3-1
RORA
PTGIS
NR1D1
CD40
NFKB1
USP10
MAPK13
DAB2IP
RELA
SFRP1
SOX9
ADAMTS12
CACTIN
ANKRD1
UPF1
ADAMTS7
CXCL8
TANK
SBNO2
CD47
ALOX15
IL18RAP
JAK2

MCM2
HSP90AB1
IMPDH2
PML
HSPA5
NFIL3
TCF7
ALAD
FASN
RUFY4
CDK4
DCSTAMP
KEAP1
CORO1A
ST3GAL6
SELPLG
PHB
PTPRT
YBX1
HDGF
RAD23B
LSP1
ATIC
BTK
GIPC1
P4HB
STIP1
EGR1
DHX9
GBA
TNFRSF21
MYOG
CHI3L1
GSDME
NFKBIA
AKT1
IKBKB
ZFAND6
RIPK1
GPER1
FABP4
CHUK
MYOD1
SLC2A4
TCL1A

CIB1
YBX3
CRHBP
ZFP36L2
HMHB1
NFE2L2
BIRC2
BAG4
MAPK1
INPP5K
SMPD3
TRADD
ERBIN
BRCA1
RPS3
ZFP36
SIRT1
ZFP36L1
GPD1
NPNT
MAPK3
IFIT1
CCL24
CCL28
CXCL12
CXCL16
CCL4L1
CXCL14
CCL2
CCL8
CCL13
CCL1
CCL5
CCL14
CCL3
CCL22
CX3CL1
CXCL13
CXCL11
CXCL10
CXCL9
CCL25
CXCL5
PPBP
PF4

CXCL1
PF4V1
CXCL6
ACKR4
XCR1
S100A14
ACKR1
PTK2B
TREM2
WBP1L
ITCH
IL5RA
CSF3R
MPL
IL12RB2
LEPR
IL6R
IFNGR1
GHR
IFNGR2
IL2RB
CSF2RB
CD74
FZD4
CD44
IL7R
IL13RA2
IL13RA1
CNTFR
IL9R
IL4R
IL21R
LIFR
OSMR
EPOR
IL6ST
PRLR
ANKRD24
IL15
IFNK
IL21
IL9
IFNA13
IFNA8
IFNA6

IFNA14
IFNA16
IFNA10
IFNW1
EDN2
GREM2
IL17RB
IRF5
IL1RAP
IL20RA
KIT
SH2B2
IFNAR1
DUOX1
IL7
SLC1A1
FER
TNFRSF1A
SOCS1
NUMBL
STAT4
SOCS5
JAK1
ZC3H15
STAT5B
JAK3
IL17RE
IL1RAPL2
IRAK3
STAT2
IL1A
IL37
IRAK4
TYK2
EREG
STAT6
TNFSF11
BAD
DLG1
DOCK2
MSN
PRF1
IFNG
IRF6
MAMU-A

IRF9
IRF4
TP53
NMI
IRF7
IL1RL2
IL1RL1
IL36RN
IL1RN
IL36G
IL36A
IL36B
IL1F10
PSMB2
PSMA5
PSMB1
PSMA6
PSMB9
PSMB8
PLCB1
PSMB5
RPS6KA5
PSMA7
PSMA1
PSMB7
PSMB4
PSMB3
IRAK1
PSMA8
IRAK2
PSMA4
RPS6KA4
IL11
IL23R
IL12A
IL15RA
SYK
CD4
IL17RD
IL17REL
TRAF5
IKBKE
NOTCH1
PDGFB
IL2RA

IL2
ECM1
IL20
IL22RA2
IL23A
IL27RA
IL27
MX1
OASL
OAS2
IFNL1
IL3
RNF41
IL33
IL34
PIBF1
IL5
SDCBP
ADAM17
ERAP1
SPI1
SRC
CTR9
YAP1
SMAD4
SHCBP1L
ECT2
MAP9
PDCD6IP
ANLN
EFHC1
APC
CUL7
SON
TRIM36
RAB35
LZTS2
CEP55
CENPA
CKAP2
SPTBN1
EFHC2
SNX9
SNX33
PLK1

RTKN
ANK3
CHMP4B
KIF20A
SNX18
MYH10
KIF4B
STAMBP
MITD1
USP8
UNC119
KIF4A
ZFYVE26
RACGAP1
CIT
BBS4
INCENP
SPAST
NUSAP1
CHMP4C
ZFYVE19
KLHDC8B
CNTROB
SLIT2
PADI2
ROBO1
SLIT3
RNF113A
PTPRC
OTOP1
PTPN2
PARP14
OTUD4
TIGIT
SLAMF1
CMKLR1
IL10
ARRB2
THBS1
C1QBP
LILRA5
TLR8
GHSR
RABGEF1
TNFAIP3

SYT11
PTPN22
HGF
NLRX1
TLR9
ORM1
PTPN6
INPP5D
NR1H4
C5AR2
NCKAP1L
BANK1
ARRB1
NLRP12
HAVCR2
NLRC3
FOXJ1
CD96
BST2
KLF4
APOA1
MAPK7
CD34
DEFB114
NFKBIL1
ILRUN
ADIPOQ
DICER1
TRIM27
RARA
VSIR
GSTP1
CIDEA
LBP
IGF1
BCL3
ARG2
CLEC4A
CD33
GPR18
MC1R
AKAP8
CD274
TRAIP
HIST1H2BJ

GPS2
XIAP
NAIP
F2RL1
PELI3
RFFL
CCDC3
NOL3
PPP2CB
PYDC1
PIAS4
MUL1
DCST1
YTHDF2
ADAR
ISG15
METTL3
SAMHD1
YTHDF3
OAS3
MMP12
TTLL12
NLRC5
TRIM32
MCOLN2
TRPV4
MAVS
ADCYAP1
LPL
MBP
CHIA
AIRE
TLR3
RIPK2
AGER
APP
ADORA2B
EIF2AK2
FFAR2
C5
TLR7
EDN1
WNT3A
GLMN
BCL10

TLR5
HHLA2
FGR
SOD1
CLEC5A
TMF1
FLOT1
BATF
BTN3A1
AGPAT1
HTR2B
PELI1
ADRA2A
LURAP1
INS
IL26
SLC11A1
CLEC9A
RAB5C
CRTAM
ITK
NFAM1
PANX1
CLEC6A
RGCC
PTGER4
CADM1
TMIGD2
CCDC88B
AGPAT2
PLA2R1
TNFRSF14
INAVA
SPHK2
HK1
LACC1
SCIMP
TRIM44
PAFAH1B1
PLCG2
DHX36
SETD2
TBK1
IRF3
DDX3X

ZBTB20
POLR3C
TOMM70
POLR3D
HMGB2
POLR3A
POLR3G
RNF135
RIOK3
DHX58
POLR3F
POLR3B
PTPN11
KCNH7
CD160
CD2
PDE4B
SLAMF6
LTA
CD244
TNFSF4
ISL1
PDE4D
CYRIB
SCRIB
NKG2D
RASGRP1
CD3E
SLC7A5
ZFPM1
HRAS
FADD
CEBPG
CD14
ABL1
HPX
MED1
PARP9
HSPB1
CD36
GSDMD
TRIM16
P2RX7
STMP1
TMEM106A

NLRP1
AZU1
USP50
CASP8
NLRC4
PANX2
PANX3
TMED10
CD46
IL20RB
TUSC2
CD40LG
CD83
CD28
LTB
MDK
IDO1
HLA-E
TIRAP
ZBTB7B
PRKCQ
OSM
TGFB1
VTCN1
RUNX1
CARD11
PRKD2
ANXA1
STOML2
CSF2
CEBPB
EPX
TLR1
ARHGEF2
NOD1
SETD4
C1QTNF4
F2R
MMP8
CYBA
POU2AF1
POU2F2
GFI1
PARK7
APOA2

SERPINE1
F3
RAB1A
GDF2
PRG3
ELANE
AFAP1L2
FCN1
DDIT3
SEMA7A
NUP62
KIF20B
DDX1
KIFBP
CLNK
CD226
ADAM8
EPHB2
TNFRSF8
NFATC4
MAPKAPK2
SPN
LY96
CLU
PSEN1
FRMD8
PIK3R1
ARFGEF2
PRKN
CASP4
TRIM15
CGAS
G3BP1
DHX33
ZCCHC3
PQBP1
TRIM56
ZBP1
LSM14A
TRIM41
USP27X
VRK2
MAST2
ANKRD53
LTF

UBE2J1
ANGPT1
NKIRAS1
HIPK1
SPATA2
OTULIN
CYLD
SHARPIN
NKIRAS2
SPHK1
ITIH4
SRF
TIMP4
MAPKAPK3
ALDH1A2
REL
BCL2L1
TMEM102
PPP3CB
AVPR2
BCL2
COL3A1
MCL1
TNFRSF11A
LAMP3
MX2
IFNAR2
SHFL
XAF1
SNCA
UBD
TRIM21
CIITA
GCH1
KYNU
CALCOCO2
CYP27B1
SP100
SELE
YTHDC2
SMPD1
IGBP1
PRKCA
SHANK3
ADAM10

MAP4K3
ADAM9
AFF3
CH25H
SHMT2
TNFSF10
SIVA1
FASLG
TRAF3
TNFSF12
BABAM2
TNFSF13B
TRIM37
TRAF1
EDA
TNFSF8
TRAF4
TNFSF15
TNFSF9
CD70
TNFSF14
TNFSF18
TNFRSF1B
FOXO3
RRAGA
ACTN4
ILK
CARD14
CDIP1
TNFRSF17
EDA2R
PLVAP
UMOD
TXNDC17
TNFRSF13C
IRF2
IFI27
