## Supplemental Table 3 for "Impact of aging on the immunological and microbial landscape of the lung during non-tuberculous mycobacterial infection"

**Supp. Table 3: The weighted intragroup Unifrac distances showing β-diversity at eac**

**Unifrac Distance BAL Right Intragroup (Weighted)**

| D0 | D8* | D15 | D28 | D44 | D56 | D86 |
|---|---|---|---|---|---|---|
| 0.092864 | 0.207105 | 0.480806 | 0.401281 | 0.450085 | 0.402182 | 0.291661 |
| 0.451782 | 0.11348 | 0.146721 | 0.148885 | 0.111169 | 0.235311 | 0.347485 |
| 0.520951 | 0.184963 | 0.522594 | 0.467354 | 0.493608 | 0.463621 | 0.432614 |
| 0.091227 | 0.133074 | 0.173406 | 0.197342 | 0.436439 | 0.293423 | 0.333313 |
| 0.004513 | 0.110487 | 0.366729 | 0.294502 | 0.028215 | 0.556618 | 0.425023 |
| 0.519961 | 0.106174 | 0.198741 | 0.232919 | 0.480212 | 0.327652 | 0.266025 |
| 0.511883 | 0.078661 | 0.251903 | 0.206874 | 0.120375 | | |
| 0.582203 | 0.222487 | 0.409119 | 0.429546 | 0.471419 | | |
| 0.320874 | 0.101572 | 0.266605 | 0.202537 | 0.120821 | | |
| 0.581478 | 0.14561 | 0.267875 | 0.187633 | 0.458576 | | |

**Unifrac Distance BAL Left Intragroup (Weighted)**

| D0 | D8 | D15 | D28 | D44 | D56 | D86 |
|---|---|---|---|---|---|---|
| 0.091485 | 0.293341 | 0.380327 | 0.384938 | 0.160687 | 0.301843 | 0.000801 |
| 0.453795 | 0.127194 | 0.255342 | 0.386843 | 0.32046 | 0.250212 | 0.31715 |
| 0.521969 | 0.256187 | 0.57019 | 0.005991 | 0.408345 | 0.438058 | 0.317411 |
| 0.089972 | 0.50422 | 0.375134 | 0.344907 | 0.258671 | 0.287076 | 0.450366 |
| 0.004823 | 0.230361 | 0.030273 | 0.458552 | 0.223255 | 0.231242 | 0.450896 |
| 0.521115 | 0.471193 | 0.562873 | 0.460698 | 0.553439 | 0.476929 | 0.465695 |
| 0.512246 | 0.167981 | 0.222765 | 0.237061 | 0.286973 | 0.297141 | 0.002634 |
| 0.580924 | 0.369947 | 0.449838 | 0.369682 | 0.354507 | 0.48374 | 0.002719 |
| 0.320577 | 0.239372 | 0.28035 | 0.370271 | 0.129508 | 0.064243 | 0.315449 |
| 0.580496 | 0.563428 | 0.444504 | 0.540121 | 0.516237 | 0.525739 | 0.448842 |

**h time point and comparing to baseline (0 DPI) for the right and left BAL**

* = p ≤ 0.05 when compared to D0

| D121 | D149 | Nx |
|---|---|---|
| 0.574553 | 0.564028 | 0.058937 |
| 0.19049 | 0.379874 | 0.480152 |
| 0.571666 | 0.516191 | 0.477061 |
| 0.279098 | 0.409453 | 0.109737 |
| 0.352816 | 0.394202 | 0.079697 |
| 0.320071 | 0.15101 | 0.510354 |
| 0.125468 | 0.317008 | |
| 0.58469 | 0.508916 | |
| 0.141294 | 0.169242 | |
| 0.293868 | 0.220529 | |

| D121 | D149 | Nx |
|---|---|---|
| 0.572637 | 0.259675 | 0.056821 |
| 0.147056 | 0.343728 | 0.482543 |
| 0.438049 | 0.473369 | 0.477219 |
| | 0.418315 | 0.108438 |
| | 0.175255 | 0.080814 |
| | 0.578431 | 0.510095 |
| | 0.150117 | |
| | 0.328243 | |
| | 0.225167 | |
| | 0.4852 | |
